# Nitrogenase resurrection and the evolution of a singular enzymatic mechanism

**DOI:** 10.1101/2022.05.17.492308

**Authors:** Amanda K. Garcia, Derek F. Harris, Alex J. Rivier, Brooke M. Carruthers, Azul Pinochet-Barros, Lance C. Seefeldt, Betül Kaçar

**Author notes:** To whom correspondence should be addressed: Betül Kaçar, [].

## Abstract

The planetary biosphere is powered by a suite of key metabolic innovations that emerged early in the history of life. However, it is unknown whether life has always followed the same set of strategies for performing these critical tasks. Today, microbes access atmospheric sources of bioessential nitrogen through the activities of just one family of enzymes, nitrogenases. Here, we show that the only dinitrogen reduction mechanism known to date is an ancient feature conserved from nitrogenase ancestors. We designed a paleomolecular engineering approach wherein ancestral nitrogenase genes were phylogenetically reconstructed and inserted into the genome of the diazotrophic bacterial model, *Azotobacter vinelandii*, enabling an integrated assessment of both *in vivo* functionality and purified nitrogenase biochemistry. Nitrogenase ancestors are active and robust to variable incorporation of one or more ancestral protein subunits. Further, we find that all ancestors exhibit the reversible enzymatic mechanism for dinitrogen reduction, specifically evidenced by hydrogen inhibition, that is also exhibited by extant *A. vinelandii* nitrogenase isozymes. Our results suggest that life may have been constrained in its sampling of protein sequence space to catalyze one of the most energetically challenging biochemical reactions in nature. The experimental framework established here is essential for probing how nitrogenase functionality has been shaped within a dynamic, cellular context to sustain a globally consequential metabolism.

**IMPACT STATEMENT:** The enzymatic mechanism for dinitrogen reduction is an ancient feature of nitrogenases that persisted over hundreds of millions of years.

## INTRODUCTION

The evolutionary history of life on Earth has generated tremendous ecosystem diversity, the sum of which is orders of magnitude larger than that which exists at present [1]. Life’s historical diversity provides a measure of its ability to solve adaptive problems within an integrated planetary system. Accessing these solutions requires a deeper understanding of the selective forces that have shaped the evolution of molecular-scale, metabolic innovations. However, the early histories of many of life’s key metabolic pathways and the enzymes that catalyze them remain coarsely resolved.

Important efforts to advance understanding of early metabolic innovations have included phylogenetic inference [2, 3], systems-level network reconstructions [4], and the leveraging of extant or mutant biological models as proxies for their ancient counterparts [5-7]. However, methods to directly study these metabolic evolutionary histories across past environmental and cellular transitions remain underexplored. A unified, experimental strategy that integrates historical changes to enzymes, which serve as the primary interface between metabolism and environment, and clarifies their impact within specific cellular and physiochemical contexts is necessary. To address this, phylogenetic reconstructions of enzymes can be directly integrated within laboratory microbial model systems [8-10]. In this paleomolecular framework, predicted ancestral enzymes can be “resurrected” within a compatible host organism for functional characterization. These experimental systems can ultimately integrate multiple levels of historical analysis by interrogating critical features of ancient enzymes as well as dynamic interactions between enzymes, their broader metabolic networks, and the external environment.

The study of biological nitrogen fixation offers a promising testbed to thread these investigations of early metabolic evolution. Both phylogenetic and geological evidence [11-15] indicate that the origin of biological nitrogen fixation was a singular and ancient evolutionary event on which the modern biosphere has since been built. The only known nitrogen fixation pathway (compared to, for example, at least seven carbon-fixation pathways [16]) is catalyzed by an early evolved family of metalloenzymes called nitrogenases that reduce highly inert, atmospheric dinitrogen (N_2_) to bioavailable ammonia (NH_3_). The nitrogenase family comprises three isozymes that vary in their metal dependence (i.e., molybdenum, vanadium, and iron) and, in certain cases, all coexist within the same host organism [17]. Many diazotrophs depend on genetic strategies for coordinating the biosynthesis and expression of multiple nitrogenase isozymes and their respective metalloclusters [18, 19], and, in oxic environments, protecting the oxygen-sensitive metalloclusters from degradation [20]. Thus, nitrogenase enzymes are a central component of a broader, co-evolving nitrogen fixation machinery. These features that create significant experimental challenges for nitrogen fixation engineering [21-23] nevertheless also make this metabolism an ideal candidate for systems-level, paleomolecular study.

How biological nitrogen fixation emerged and evolved under past environmental conditions is still poorly constrained relative to its importance in Earth’s planetary and biological history. Because nitrogen has been a limiting nutrient over geological timescales [24, 25], nitrogenase has long been a key constituent of the expanding Earth biosphere. The impact of nitrogen limitation is underscored by human reliance on the industrial Haber-Bosch process, an energetically and environmentally costly workaround for nitrogen fertilizer production [26] designed to supplement a remarkable molecular innovation that biology has tinkered with for more than three billion years. How the structural domains and regulatory network of nitrogenase were recruited [27, 28] and under what selective pressures the metal dependence of nitrogenases were shaped [11, 29] remain open questions. Importantly, it is not known how the enzymatic mechanism for dinitrogen reduction has been tuned by both peptide and metallocluster to achieve one of the most difficult reactions in nature [30-32]. At the enzyme level, previous insights into nitrogenase sequence-function relationships have primarily derived from single or dual substitution studies. These have often yielded diminished or abolished nitrogenase activity [30, 31], though in certain cases improved reactivity toward alternate, industrially relevant substrates [30]. Despite illuminating key features of extant nitrogenase mechanism in select model organisms, the combination of detailed functional studies within an explicit evolutionary scheme has not previously been accomplished for the nitrogen fixation system.

Here, we seek guidance from the Earth’s evolutionary past to reconstruct the history of the key metabolic enzyme, nitrogenase. We establish an evolutionary systems biology approach for the cellular- and molecular-level characterization of ancestral nitrogenases resurrected within the model diazotrophic bacterium, *Azotobacter vinelandii* (*A. vinelandii*). We find that variably replacing different protein subunits of the nitrogenase complex with inferred ancestral counterparts enables nitrogen fixation in *A. vinelandii*. Purified ancestral enzymes exhibit the specific N_2_ reduction mechanism retained by their studied, extant counterparts, and maintain the same catalytic selectivity between N_2_ and protons. Thus, the core strategy for biological nitrogen fixation is conserved across the investigated timeline. Our paleomolecular approach opens a new route to study the ancient functionality and evolution of nitrogenases both deeper in its ancestry and within the broader context of its supporting, cellular machinery.

## RESULTS

### A resurrection strategy for ancestral nitrogenases

We designed an engineering pipeline for the resurrection and experimental characterization of ancestral nitrogenases (**Fig. 1*A***). In this scheme, phylogenetically inferred, ancestral nitrogenase genes are synthesized and engineered into the genome of a modern diazotrophic bacterium, enabling the assessment of *in vivo* nitrogenase activity and expression in parallel with biochemical analysis of purified enzyme. The engineering and functional assessment of ancestral nitrogenases required a suitable diazotrophic microbial host, owing to challenges associated with nitrogenase heterologous expression [26]. We selected the obligately aerobic gammaproteobacterium, *A. vinelandii* (strain “DJ”), an ideal experimental model due to its genetic tractability and the availability of detailed studies on the genetics and biochemistry of its nitrogen fixation machinery [33]. We specifically targeted the extant *A. vinelandii* molybdenum-dependent (“Mo-”) nitrogenase (hereafter referred to as wild-type, “WT”), which is the best studied isozyme [30] relative to the *A. vinelandii* vanadium (“V-”) and iron (“Fe-”) dependent nitrogenases. The WT Mo-nitrogenase complex comprises multiple subunits, NifH, NifD, and NifK (encoded by *nifHDK* genes), which are arranged into two catalytic components: a NifH homodimer and a NifDK heterotetramer (**Fig. 1*B***). During catalysis, both components transiently associate to transfer one electron from NifH to NifDK and subsequently dissociate. Transferred electrons accumulate at the active-site Mo-containing metallocluster (“FeMoco”) housed within the NifD subunits for reduction of N_2_ substrate to NH_3_.

**Fig. 1.**
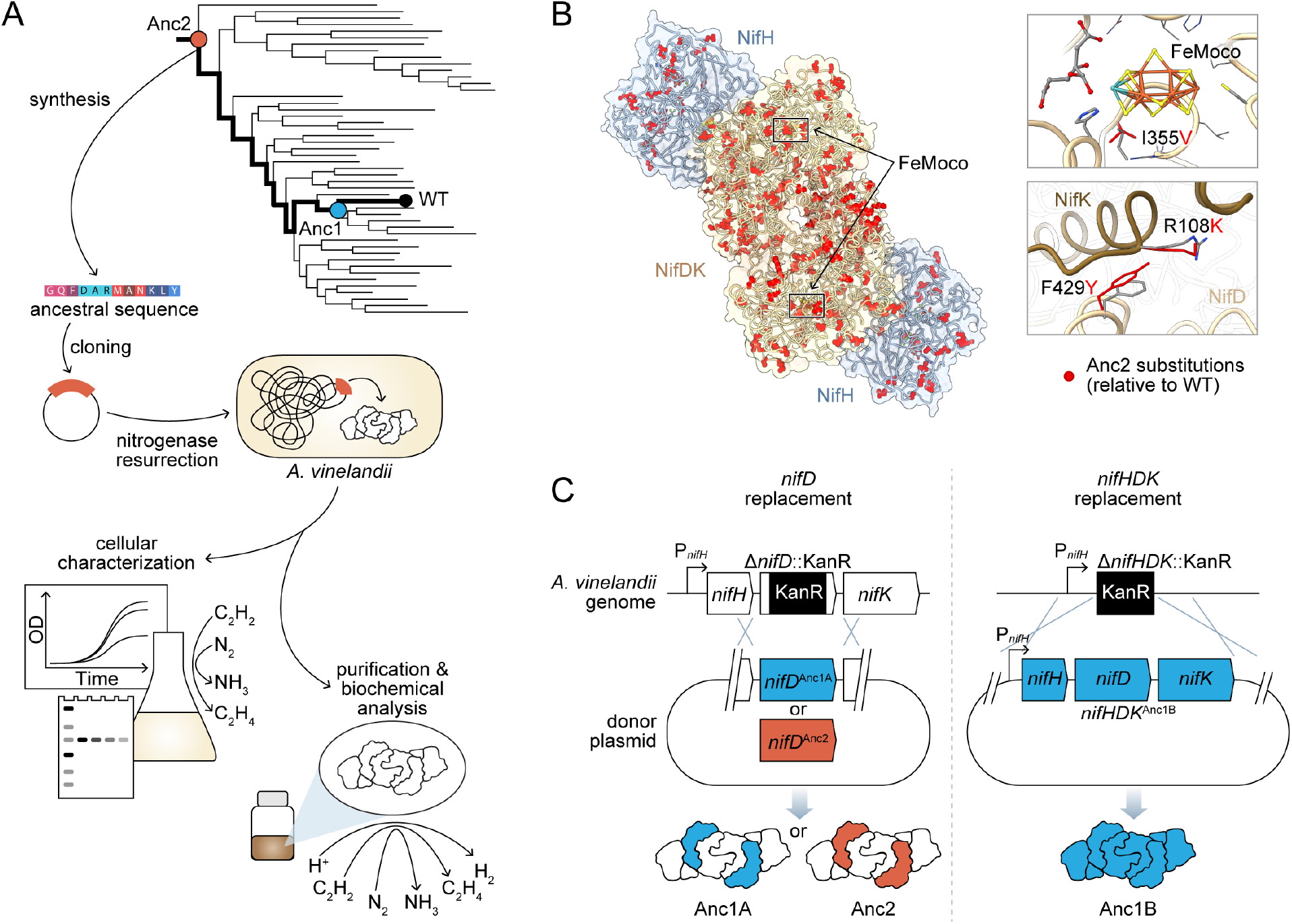
Engineering strategy for ancestral nitrogenase resurrection. (*A*) Experimental pipeline for nitrogenase resurrection in *A. vinelandii* and subsequent characterization, as described in the main text. (*B*) Structural overview of ancestral nitrogenases reconstructed in this study. Homology models (template PDB 1M34) of Anc1B NifH and NifDK proteins are shown with ancestral substitutions (relative to WT) highlighted in red. Select substitutions at relatively conserved sites in proximity to FeMoco (NifD, I355V) and the NifD:NifK interface (NifD, F429Y; NifK, R108K) are displayed in the insets. (*C*) Parallel genome engineering strategies executed in this study, involving both ancestral replacement of only *nifD* (Anc1A and Anc2) and replacement of *nifHDK* (Anc1B). “P_*nifH*_”: *nifH* promoter, “KanR”: kanamycin resistance cassette. *Anc1A and Anc1B were each reconstructed from equivalent nodes of alternate phylogenies (see **Materials and methods**).

To infer Mo-nitrogenase ancestors, we built a maximum-likelihood nitrogenase phylogeny from a concatenated alignment of NifHDK amino acid sequences (**Fig. 2*A*; Fig. 2-Figure supplement 1**). The phylogeny contains 385 sets of homologs representative of known nitrogenase molecular sequence diversity (including Mo-, V-, and Fe-nitrogenases), and is rooted by dark-operative protochlorophyllide oxidoreductase proteins classified within the nitrogenase superfamily [34]. For this study, we selected ancestors that fall within the direct evolutionary lineage of *A. vinelandii* WT (**Fig. 2*B***), “Anc1” and “Anc2” (listed in order of increasing age), having ∼90% and ∼85% amino acid sequence identity to WT across the full length of their concatenated NifHDK proteins, respectively (**Fig. 2*C*; Supplementary File 1a**). A relatively conservative percentage identity threshold was chosen based on prior studies benchmarking functional expression of ancestral elongation factor proteins in *Escherichia coli* [35] and *Synechococcus elongatus* [10]. The high-dimensional, nitrogenase protein sequence space occupied by both extant and ancestral homologs is visualized in two-dimensions in **Fig. 2*D*** by machine-learning embeddings (see **Materials and methods**). This analysis highlights the swath of sequence space targeted here, as well as that made accessible by resurrection of nitrogenase ancestors more broadly. WT, Anc1, and Anc2 lie within a Mo-nitrogenase clade (previously termed “Group I” [12]) that contains homologs from diverse aerobic and facultatively anaerobic taxa, including proteobacteria and cyanobacteria (**Fig. 2*A***). A maximum age constraint of ∼2.5 Ga for Group I nitrogenases (and thus for both Anc1 and Anc2) can be reasoned based on timing of the Great Oxidation Event [36] and downstream emergence of aerobic taxa represented nearly exclusively within this clade.

**Fig. 2.**
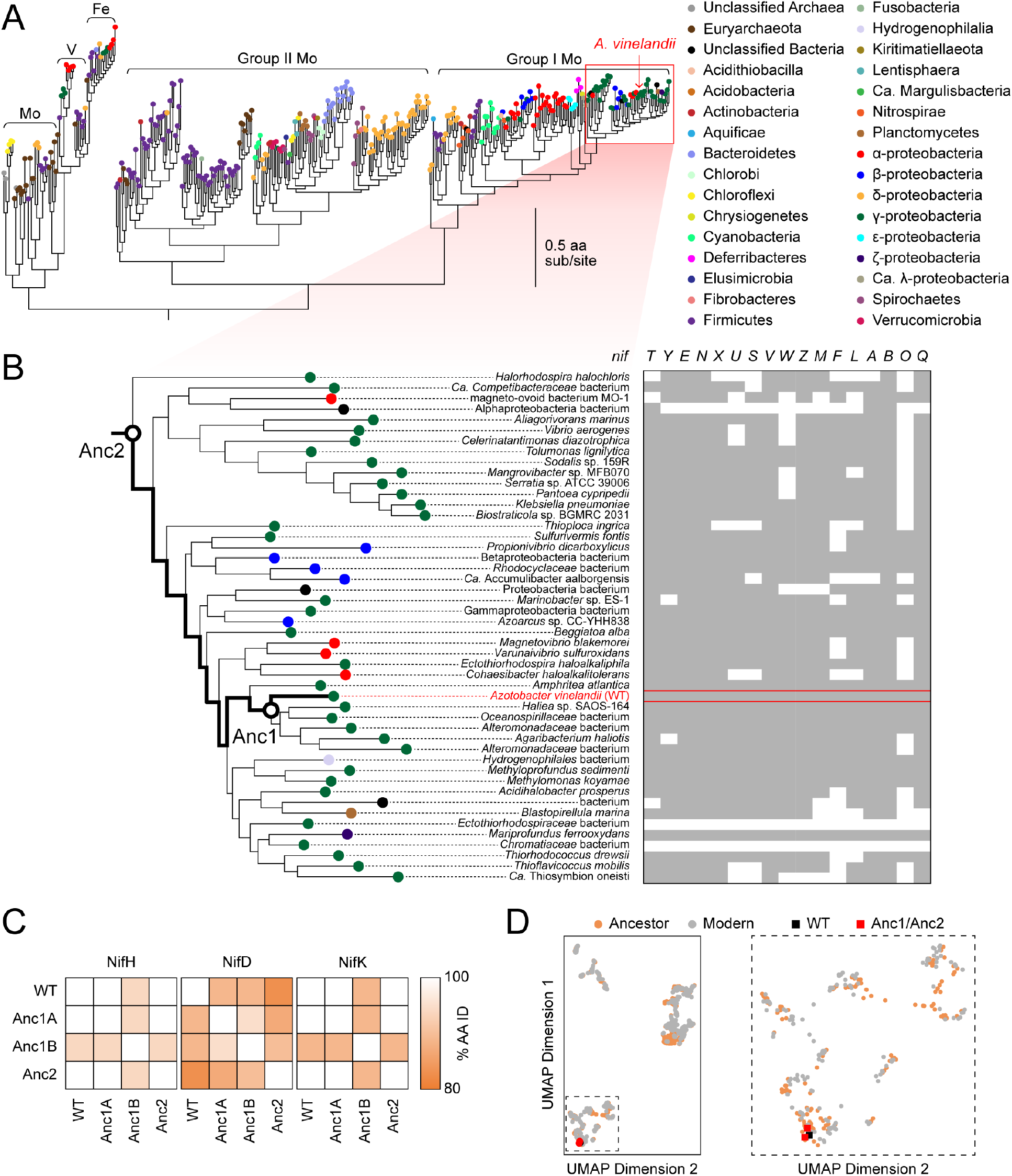
Phylogenetic and genomic context of resurrected ancestral nitrogenases. (*A*) Maximum-likelihood phylogenetic tree from which ancestral nitrogenases were inferred. Extant nodes are colored by microbial host taxonomic diversity. Red box highlights the clade targeted in this study and depicted in (*B*). Tree shown was used to infer Anc1A and Anc2 sequences (an alternate tree was used for Anc1B inference; see **Materials and methods**). (*B*) *nif* gene cluster complexity within the targeted nitrogenase clade. Presence and absence of each *nif* gene are indicated by grey and white colors, respectively. Because some homologs for phylogenetic analysis were obtained from organisms lacking fully assembled genomes, the absence of accessory *nif* genes may result from missing genomic information. (*C*) Amino acid sequence identity matrix of nitrogenase protein subunits harbored by WT and engineered *A. vinelandii* strains. (*D*) Extant and ancestral nitrogenase protein sequence space visualized by machine-learning embeddings, with the resulting dimensionality reduced to two-dimensional space. UMAP dimension axes are in arbitrary units. The field demarcated by dashed lines in the left plot is expanded on the right plot.

Residue-level differences between ancestral and WT nitrogenases (“ancestral substitutions”) are broadly distributed along the length of each ancestral sequence (**Fig. 1*B*; Fig. 2-Figure Supplement 2**). An ancestral substitution proximal to the active-site FeMoco metallocluster lies within a loop considered important for FeMoco insertion [37] (NifD I355V; residue numbering from WT)) and is observed across all targeted NifD ancestors (**Fig. 1*B***). Other ancestral substitutions are notable for their location at relatively conserved residue sites (assessed by ConSurf [38]; **Fig. 2-Figure Supplement 3**; see **Materials and methods**) and/or within subunit interfaces, including two at the NifD:NifK interface that are proximal to one another, F429Y (NifD) and R108K (NifK). The Anc2 NifD protein contains five more ancestral substitutions at conserved sites than the younger Anc1 NifD protein. In all studied ancestors, the C275 and H442 FeMoco ligands, as well as other strictly conserved nitrogenase residues, are retained.

Phylogenetic analysis informs the compatibility of selected ancestors in extant microbial hosts. Extant nitrogenases within the Group I nitrogenase clade (which include Anc1 and Anc2 descendants) are associated with numerous accessory genes likely recruited to optimize synthesis and regulation of the oxygen-sensitive nitrogenase for aerobic or facultative metabolisms [27]. For example, in addition to the structural *nifHDK* genes, *A. vinelandii* WT is assembled and regulated with the help of >15 additional *nif* genes. Likewise, the extant descendants of Anc1 and Anc2 are primarily aerobic or facultative proteobacteria and are thus associated with higher complexity *nif* gene clusters (**Fig. 2*B***). We hypothesized that the likely oxygen-tolerant, ancient proteobacterium harboring these ancestral nitrogenases were similar in *nif* cluster complexity to extant *A. vinelandii*. Thus, we predicted that the *nif* accessory genes present in *A. vinelandii* would support the functional expression of resurrected nitrogenase ancestors.

To gauge the degree of compatibility between ancestral and extant nitrogenase proteins and maximize the chance of recovering functional nitrogenase variants, we executed two parallel genome engineering strategies. First, we constructed *A. vinelandii* strains harboring only the ancestral *nifD* gene from both targeted ancestral nodes (“Anc1A” and “Anc2”), thereby expressing “hybrid ancestral-WT” nitrogenase complexes (**Fig. 1*C***). This strategy is similar to *in vitro* “cross-reaction” studies that have evaluated the compatibility of nitrogenase protein components from differing host taxa [39]. Second, we constructed a strain harboring all Anc1 *nifHDK* genes, expressing a fully ancestral nitrogenase complex (“Anc1B”; sequence reconstructed from a node equivalent to Anc1A from an alternate phylogeny, see **Materials and methods**). *A. vinelandii* strains were constructed by markerless genomic integration of ancestral nitrogenase genes, as described in **Materials and methods**.

### Ancestral nitrogenases enable diazotrophic microbial growth

All *A. vinelandii* constructs harboring ancestral genes enabled diazotrophic growth in molybdenum-containing, nitrogen-free media. All strains had comparable doubling times to WT during the exponential phase (*p* > .05; **Fig. 3*A*, 3*B***). The only significant difference among strains was a ∼14-h increase in the lag phase of strain Anc2 relative to WT, harboring the oldest nitrogenase ancestor (*p* ≈ 2e-7). We did not detect growth under the same conditions for a control Δ*nifD* strain (DJ2278, see **Supplementary File 1c**). This result confirmed that the growth observed for ancestral strains did not stem from leaky expression of the alternative, V- or Fe-dependent nitrogen fixation genes in *A. vinelandii*, which were left intact.

**Fig. 3.**
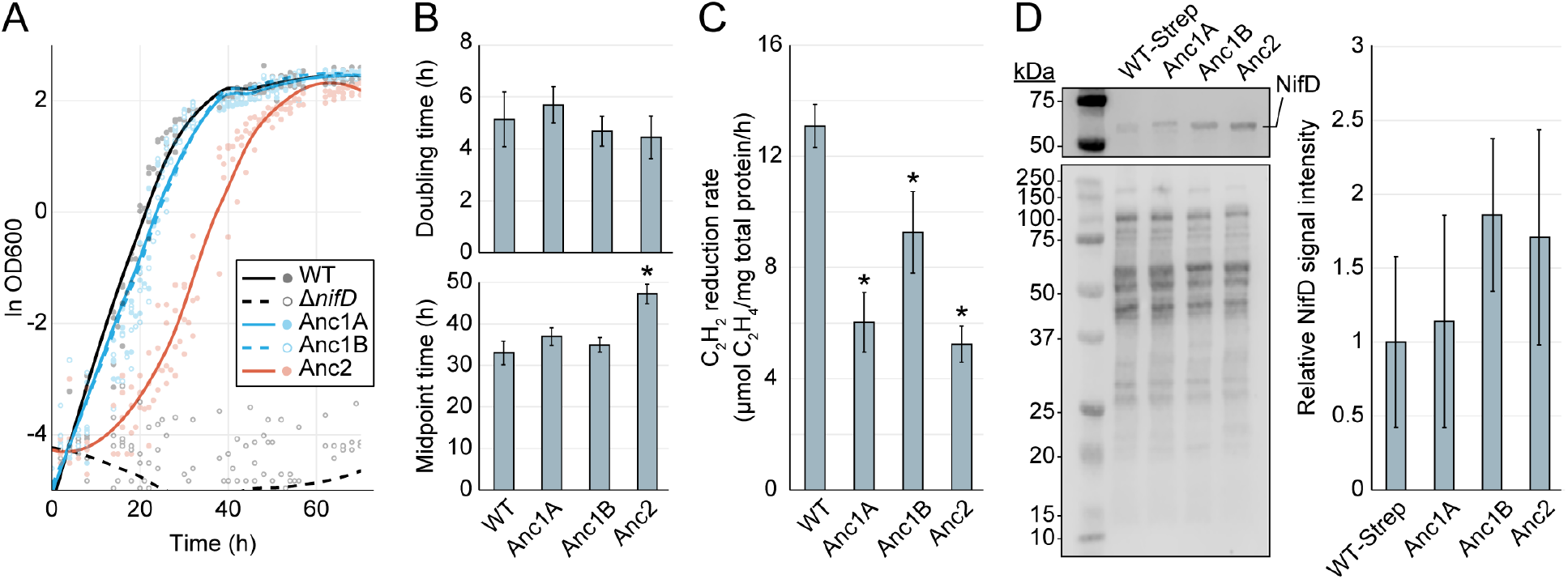
Cellular-level characterization of ancestral nitrogenase activity and expression. (*A*) Diazotrophic growth curves of *A. vinelandii* strains measured by the optical density at 600 nm (“OD600”). A smoothed curve is shown alongside individual data points obtained from five biological replicates per strain. The non-diazotrophic DJ2278 (Δ*nifD*) strain was used as a negative control. (*B*) Mean doubling and midpoint times of *A. vinelandii* strains, calculated from data in (*A*). (*C*) *In vivo* acetylene (C_2_H_2_) reduction rates quantified by production of ethylene (C_2_H_4_). Bars represent the mean of three biological replicates per strain. (*D*) Immunodetection and protein quantification of Strep-II-tagged WT (“WT-Strep”, strain DJ2102) and ancestral NifD. Top gel image shows Strep-II-tagged NifD proteins detected by anti-Strep antibody and bottom gel image shows total protein stain. Plot displays relative immunodetected NifD signal intensity normalized to total protein intensity and expressed relative to WT. Bars in plot represent the mean of three biological replicates per strain. (*B-D*) Error bars indicate ± 1 SD and asterisks indicate *p* < .01 (one-way ANOVA, post-hoc Tukey HSD) compared to WT or WT-Strep.

An acetylene reduction assay was performed to measure cellular nitrogenase activity in engineered strains. This assay quantifies the reduction rate of the non-physiological substrate acetylene (C_2_H_2_) to ethylene (C_2_H_4_) [40], here normalized to total protein content. *A. vinelandii* strains harboring only ancestral *nifD* (Anc1A, Anc2) exhibited mean C_2_H_2_ reduction rates of ∼5 to 6 μmol C_2_H_4_/mg total protein/h, ∼40-45% that of WT (*p* ≈ 6e-4 and *p* ≈ 3e-4, respectively) (**Fig. 3*C***). Strain Anc1B, harboring ancestral *nifHDK*, exhibited a mean acetylene reduction rate of ∼9 μmol C_2_H_4_/mg total protein/h, ∼70% that of WT (*p* ≈ 3e-2).

The phenotypic variability we observed among engineered and WT *A. vinelandii* strains might result both from differences in nitrogenase expression and nitrogenase activity. To provide insights into these disparate effects, we quantified nitrogenase protein expression in engineered strains by immunodetection of ancestral and WT Strep-tagged NifD proteins (the latter from strain DJ2102, see **Supplementary File 1c**) and did not conclusively detect significant differences in protein quantity relative to WT (*p* > .05; **Fig. 3*D***).

### Purified ancestral nitrogenases conserve extant N_2_ reduction mechanism and efficiency

Ancestral nitrogenase NifDK protein components were expressed and purified for biochemical characterization. All ancestral NifDK proteins were assayed with WT NifH protein (ancestral NifH proteins were not purified) for reduction of H^+^, N_2_, and C_2_H_2_. Ancestors were found to reduce all three substrates *in vitro*, supporting the cellular-level evidence of ancestral nitrogenase activity (**Fig. 4*A***).

**Fig. 4.**
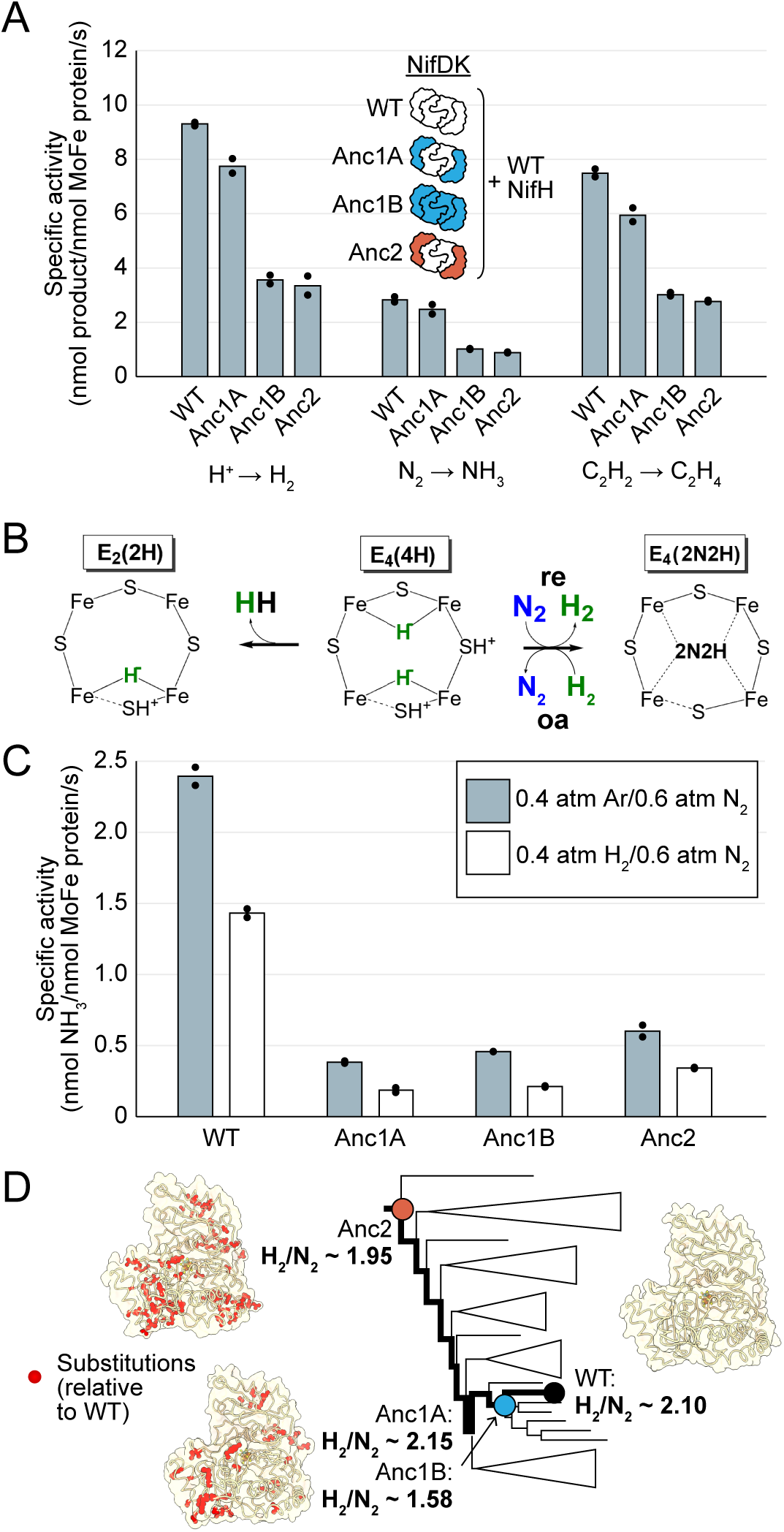
*In vitro* analyses of ancestral nitrogenase activity profiles and mechanism. All measurements obtained from assays using purified NifDK assayed with WT NifH. (*A*) Specific activities measured for H^+^, N_2_, and C_2_H_2_ substrate. (*B*) Partial schematic of the reductive-elimination N_2_-reduction mechanism of nitrogenase is shown above, centering on the N_2_-binding E_4_(4H) state of FeMoco (see main text for discussion) [32]. (*C*) Inhibition of N_2_ reduction by H_2_, evidencing the mechanism illustrated in (*B*). (*D*) Catalytic efficiencies of ancestral nitrogenases, described by the ratio of formed H_2_ to reduced N_2_ (H_2_/N_2_), mapped across the targeted phylogenetic clade. NifD homology models (PDB 1M34 template) are displayed with ancestral substitutions highlighted in red. (*A,C*) Bars represent the mean of two independent experiments with individual data points shown as black circles.

We investigated whether the ancestral nitrogenases studied here would exhibit the general mechanism for N_2_ binding and reduction that has been observed for the studied, extant nitrogenase isozymes of *A. vinelandii* (Mo, V, and Fe) [32, 41]. This mechanism involves the accumulation of 4 electron/protons on the active-site cofactor as metal-bound hydrides, generating the E_4_(4H) state (**Fig. 4*B***). Once generated, N_2_ can bind to the E_4_(4H) state through a reversible reductive elimination/oxidative addition (*re*/*oa*) mechanism, which results in the release (*re*) of a single molecule of hydrogen gas (H_2_). N_2_ binding is reversible in the presence of sufficient H_2_, which displaces bound N_2_ and results in reformation of E_4_(4H) with two hydrides (*oa*). Thus, a classic test of the (*re*/*oa*) mechanism is the ability of H_2_ to inhibit N_2_ reduction. We observed that the reduction of N_2_ to NH_3_ for all nitrogenase ancestors was inhibited in the presence of H_2_, indicating that the ancestors follow the same mechanism of N_2_ binding determined for extant enzymes (**Fig. 4*C***).

In the event the E_4_(4H) state fails to capture N_2_, nitrogenases will simply produce H_2_ from the E_4_(4H) state to generate the E_2_(2H) state. The ratio of H_2_ formed to N_2_ reduced (H_2_/N_2_) can be used as a measure of the efficiency of nitrogenases in using ATP and reducing equivalents for N_2_ reduction. The stoichiometric minimum of the mechanism is H_2_/N_2_ = 1. Experimentally (under 1 atm N_2_), a ratio of ∼2 is seen for Mo-nitrogenase and ∼5 and ∼7 for V- and Fe-nitrogenase, respectively [32]. H2/N_2_ values for all ancestors under 1 atm N_2_ was ∼2, similar to extant Mo-nitrogenase (**Fig. 4*D***).

## DISCUSSION

In this study, we leverage a new approach to investigate ancient nitrogen fixation by the resurrection and functional assessment of ancestral nitrogenase enzymes. We demonstrate that engineered *A. vinelandii* cells can reduce N_2_ and C_2_H_2_ and exhibit diazotrophic growth rates comparable to WT, though we observe that the oldest ancestor, Anc2, has a significantly longer lag phase. Purified nitrogenase ancestors are active for reduction of H^+^, N_2_, and C_2_H_2_, while maintaining the catalytic efficiency (described by the H_2_/N_2_ ratio) of WT enzymes. Our results also show that ancestral N_2_ reduction is inhibited by H_2_, indicating an early emergence of the reductive-elimination N_2_-reduction mechanism preserved by characterized, extant nitrogenases [32, 41]. These properties are maintained despite substantial residue-level changes to the peripheral nitrogenase structure (including relatively conserved sites), as well as a handful within the active-site or protein-interface regions within the enzyme complex.

It is important to consider that the nitrogenase ancestors resurrected here represent hypotheses regarding the true ancestral state. Uncertainty underlying ancestral reconstructions might derive from incomplete extant molecular sequence data, as well as incorrect assumptions associated with the implemented evolutionary models [8]. For example, a complicating feature of nitrogenase evolution is that it has been shaped significantly by horizontal gene transfer [12, 15], which in certain cases has led to different evolutionary trajectories of individual nitrogenase structural genes. Specifically, certain H-subunit genes of the V-nitrogenase appear to have different evolutionary histories relative to their DK components [12]. However, since our study targets a Mo-nitrogenase lineage, we do not expect horizontal transfer to be a significant source of uncertainty in our reconstructions.

The N_2_-reduction activity of nitrogenase ancestors suggests that the required protein-protein interactions—both between subunits that comprise the nitrogenase complex as well as those required for nitrogenase assembly in *A. vinelandii*—and metallocluster interactions are sufficiently maintained for primary function. Still, our results reveal the degree to which the organism-level phenotype of host strains can be perturbed by varying both the number and age of ancestral subunits. Importantly, these changes appear to impact phenotypic properties in complex ways, representative of the type of cellular constraints on nitrogenase evolution that would be unobservable through an *in vitro* study alone. For example, we observed comparable growth characteristics of strains harboring single (Anc1A, ancestral NifD) versus multiple (Anc1B, ancestral NifHDK) ancestral subunits of equivalent age, whereas the lag phase of Anc2 hosting a single, older subunit (ancestral NifD) was increased. Here, growth is sensitive to older, ancestral substitutions in a single subunit while permissive of more recent ancestral substitutions across one or more subunits within the nitrogenase complex. However, a different pattern is observed across *in vivo* acetylene reduction rates. These are most negatively impacted relative to WT in strains with a single NifD ancestor (Anc1A, Anc2), whereas rates are more modestly decreased in a strain with a complete, ancestral NifHDK complex (Anc1A). These results suggest that ancestral subunits of equivalent age have greater compatibility and yield greater *in vivo* activity compared to subunits of disparate ages, perhaps owing to modified protein interactions within the nitrogenase complexes The discrepancy between *in vivo* activity and growth characteristics may also be attributable to impacted cellular processes external to the biochemical properties of the nitrogenase complex itself, and yet nevertheless vital in determining the overall fitness of the host organism. Finally, though we do not detect significant differences in ancestral protein expression here, it is possible that phenotypic outcomes of future reconstructions might be impacted by perturbed expression levels (e.g., [10, 42]). To what degree these expression levels are representative of the ancestral state and impact the phenotypic property of interest should be considered in future work.

That nitrogenase ancestors perform the reductive-elimination N_2_-reduction mechanism—as the distantly related [11], extant Mo-, V-, and Fe-nitrogenases of *A. vinelandii* do today [32]—likely indicates that this enzymatic characteristic was set early in nitrogenase evolutionary history and sustained through significant past environmental change [36, 43, 44] and ecological diversification [27, 45]. It is possible that life’s available strategies for achieving N_2_ reduction may be fundamentally limited, and that a defining constraint of nitrogenase evolution has been the preservation of the same N_2_ reduction mechanism across shifting selective pressures. For example, in the acquisition of V- and Fe-dependence from Mo-dependent ancestors [11], nitrogenases may have required substantial sequence and structural changes [46, 47] in order to facilitate reductive-elimination given a different active-site metallocluster. It is also possible that alternate strategies for biological nitrogen fixation evolved early in the history of life and were subsequently outcompeted, leaving no trace of their existence in extant microbial genomes. Why these alternate possibilities were evidently not explored by nature to the same degree remains an open question, particularly given the several abiotic mechanisms for nitrogen fixation [48-50] and the multiple biological pathways for another, globally significant metabolism, carbon fixation [16]. Because our paleomolecular approach is ultimately informed by extant sequence data, it cannot directly evaluate extinct sequences that, for instance, due to contingency or entrenchment, did not persist and become preserved in extant microbial genomes. Nevertheless, evolutionarily informed studies of nitrogenase functionality that define the sequence-function space of this enzyme family will provide a foundation for laboratory efforts aimed toward exploring alternate scenarios. Future work that explores deeper into nitrogenase evolutionary history (and across extant and ancestral nitrogenase sequence space, as charted here (**Fig. 2*D***)) will clarify the degree of functional constraint exhibited by the nitrogenase family, both past and present.

## CONCLUSION

Broadening the historical level of analysis beyond a single enzyme to the organism level is necessary to generate comprehensive insights into the evolutionary history and engineering potential of nitrogen fixation. Paleomolecular work that has expanded toward the systems-level investigation of early evolved, crucial metabolic pathways remains in its infancy, despite the potential for provocative connections between molecular-scale innovations and planetary history [8, 10]. Our results highlight the evolutionary conservation of a critical metabolic pathway that has shaped the biosphere over billions of years, as well as establish the tractability of leveraging phylogenetic models to carry out extensive, empirical manipulations of challenging enzymatic systems and their microbial hosts. Building on the empirical framework presented here will illuminate the evolutionary design principles behind ancient metabolic systems more broadly as well as leverage these histories to understand how key enzymes that allowed organisms to access nitrogen from the atmosphere evolved.

## MATERIALS AND METHODS

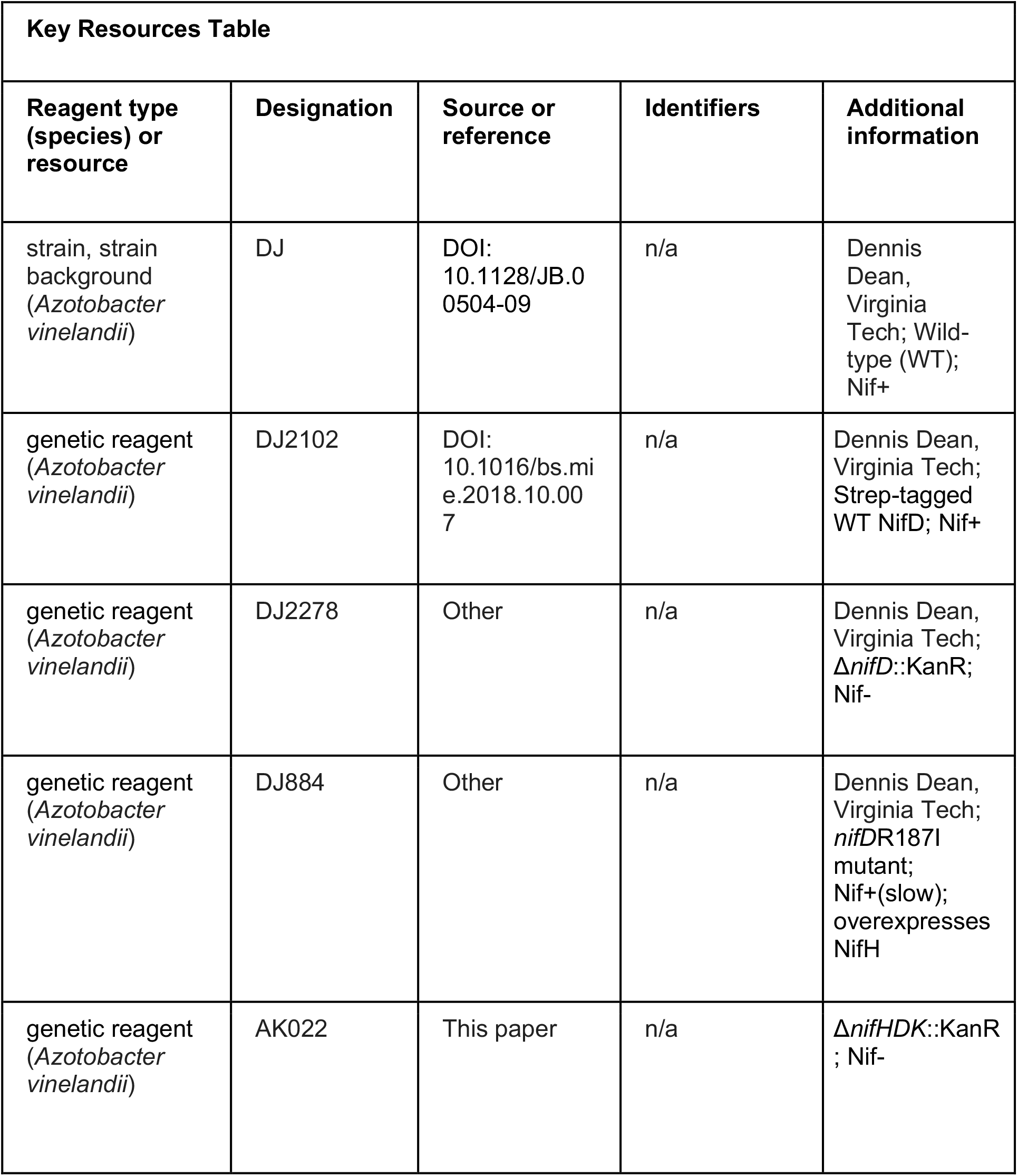
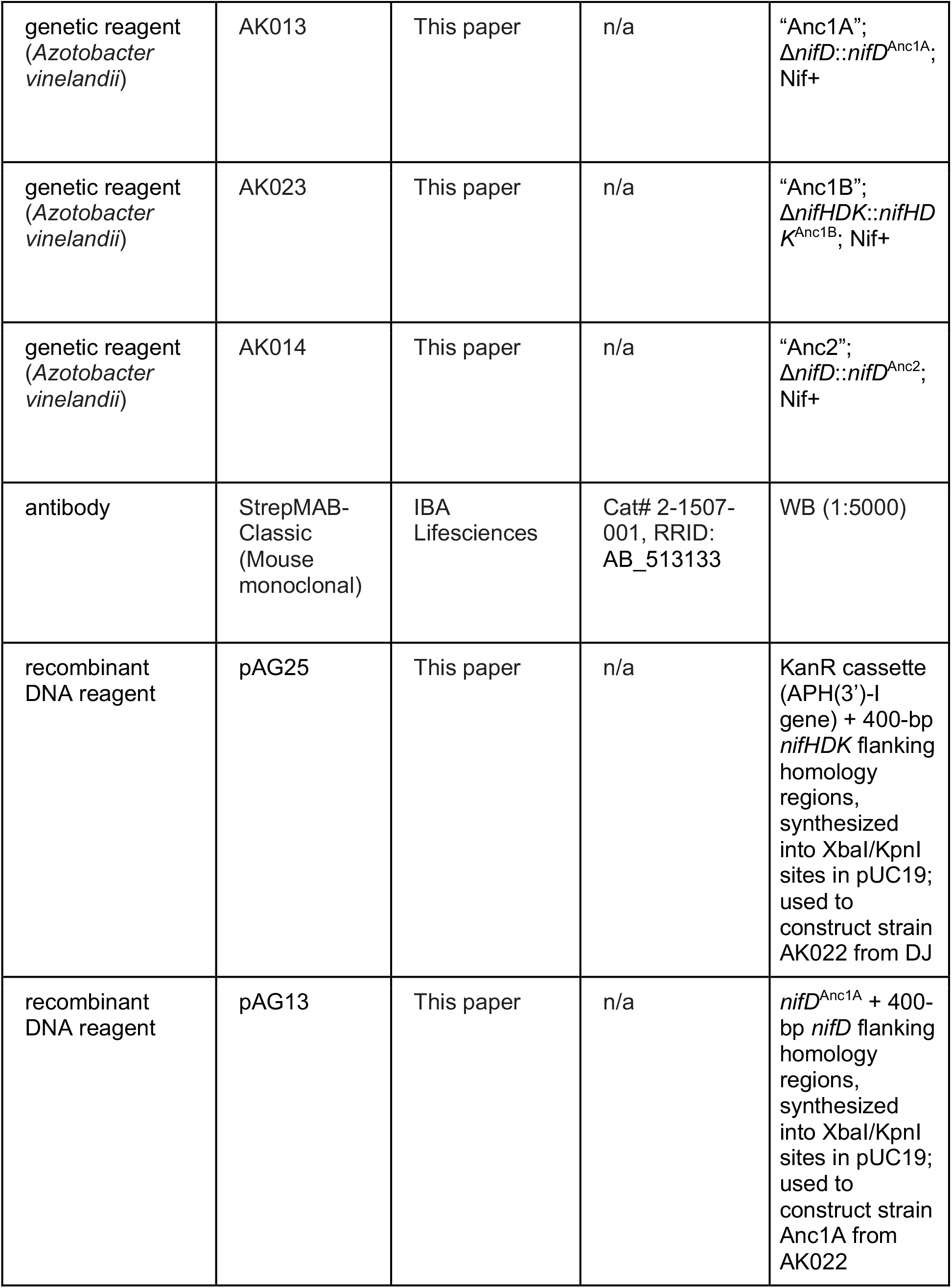
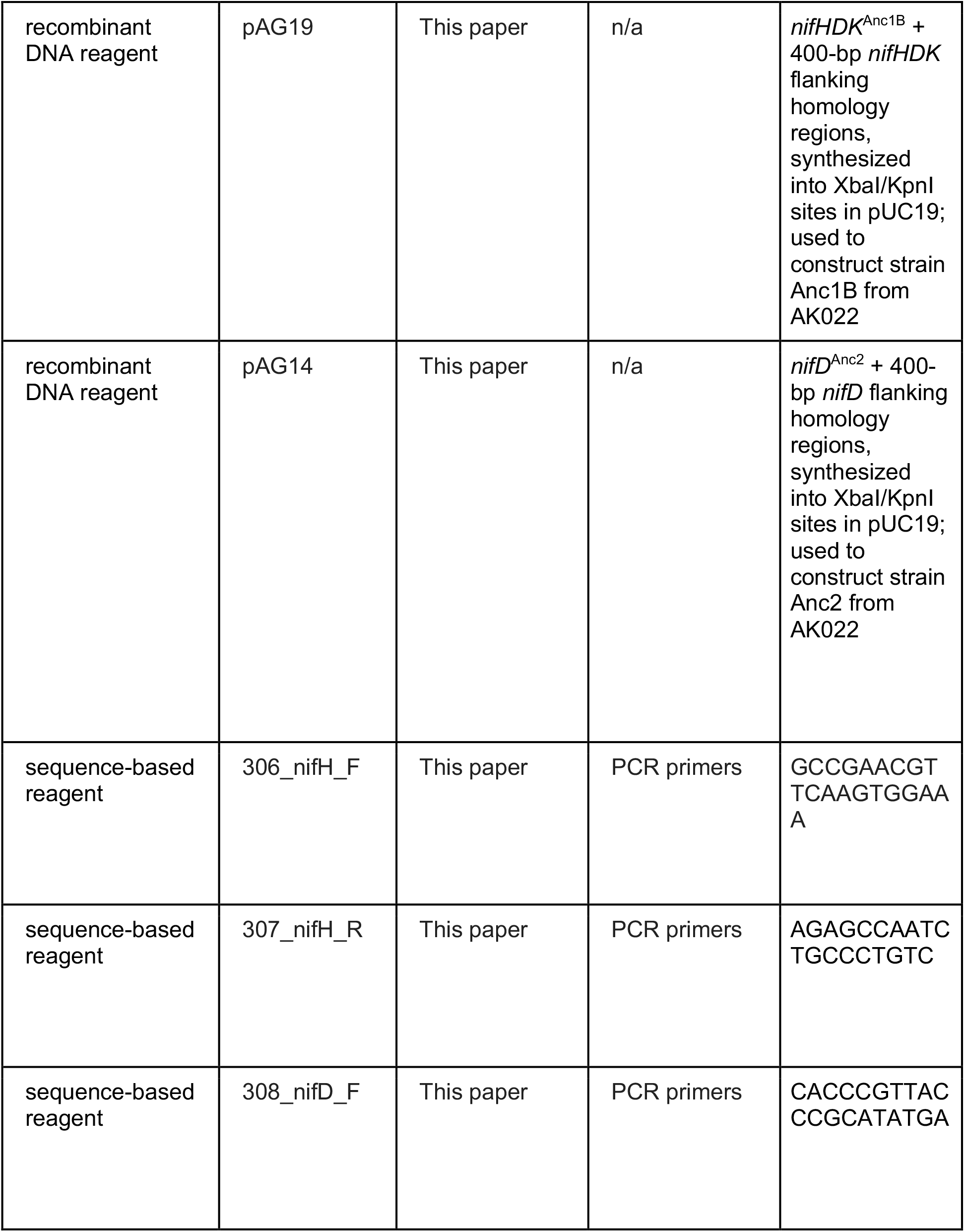
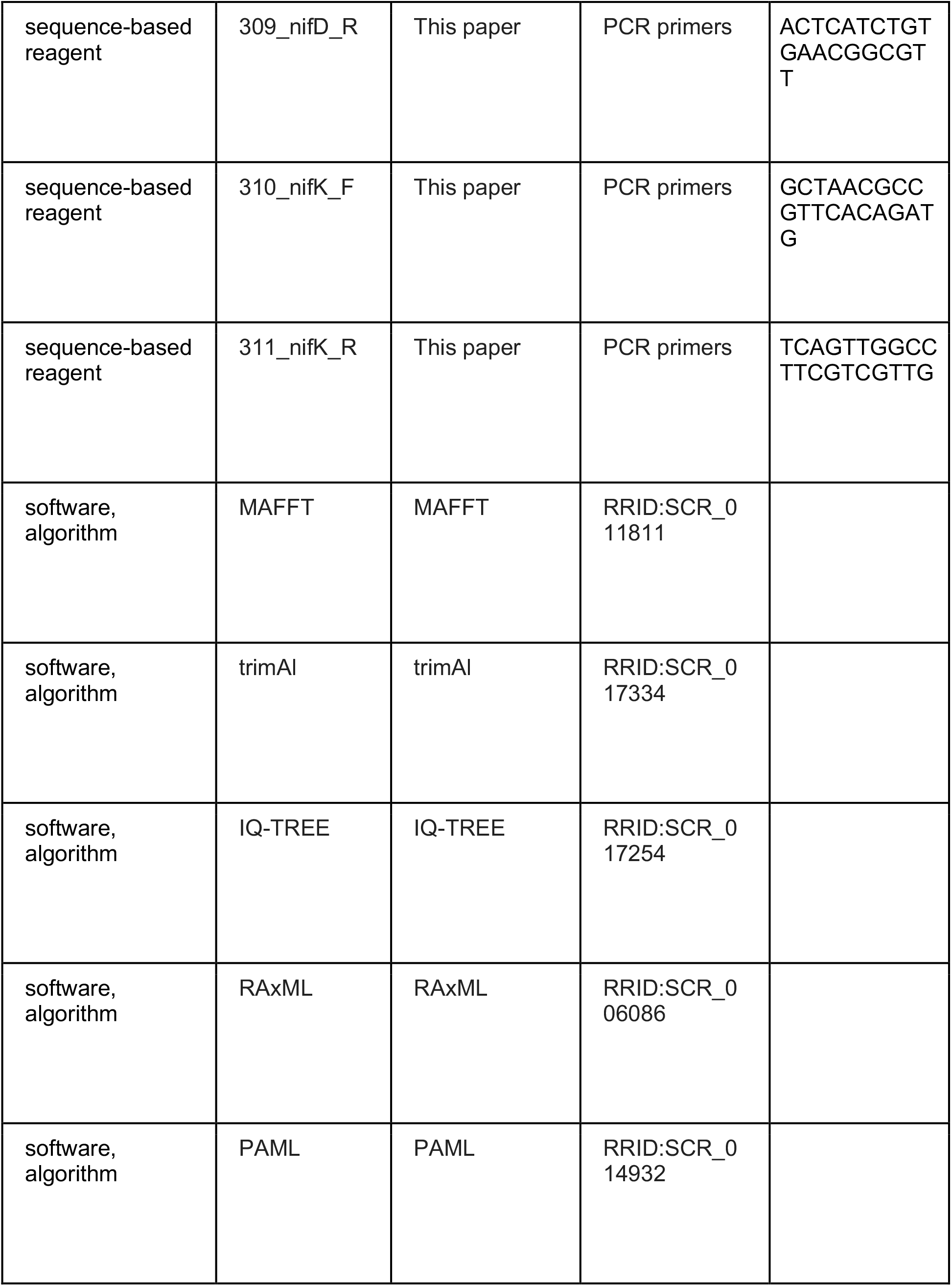
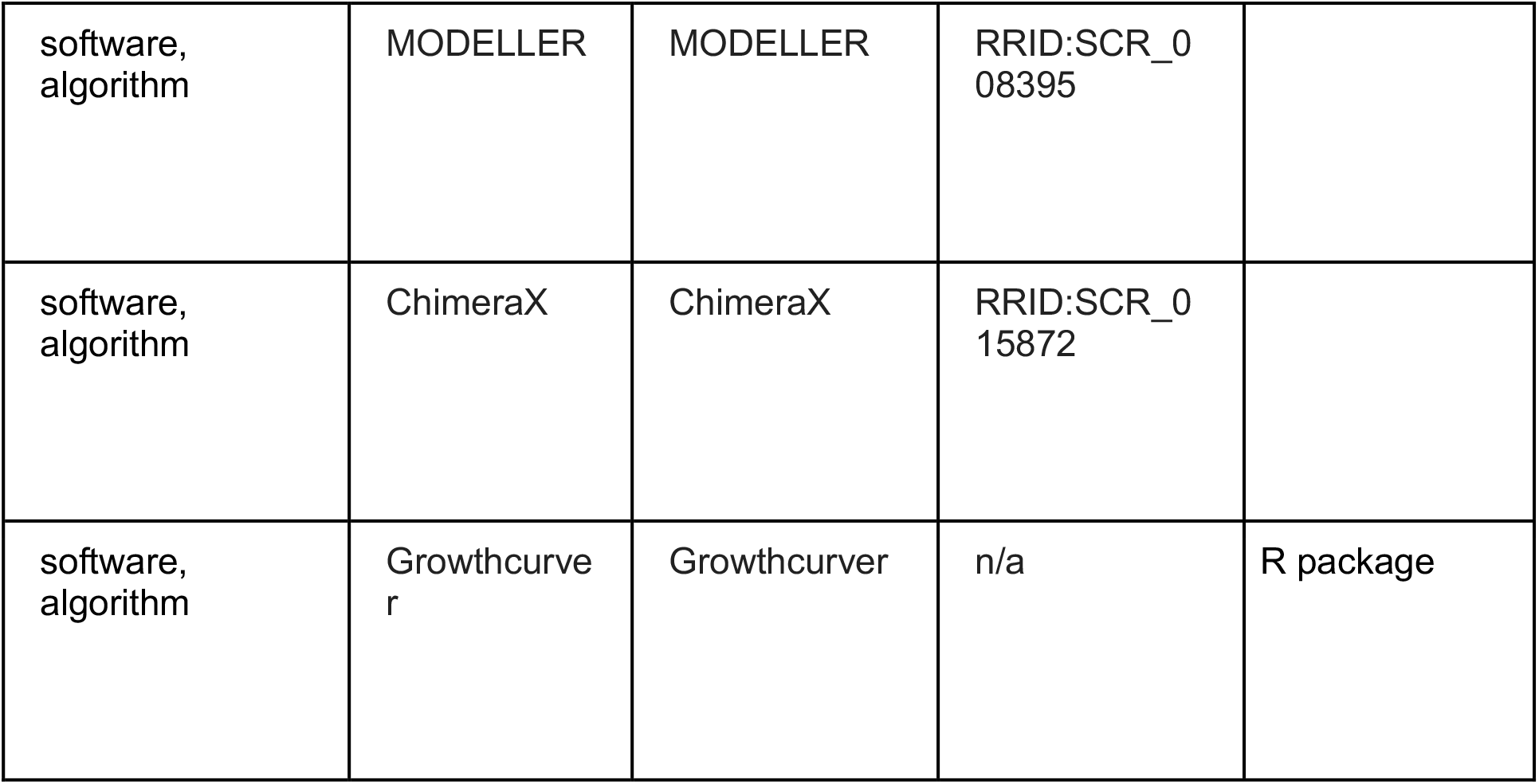

### Nitrogenase ancestral sequence reconstruction and selection

The nitrogenase protein sequence dataset was assembled by BLASTp [51] search of the NCBI non-redundant protein database (accessed August 2020) with *A. vinelandii* NifH (WP_012698831.1), NifD (WP_012698832.1), and NifK (WP_012698833.1) queries and a 1e-5 Expect value threshold (**Supplementary File 1b**). BLASTp hits were manually curated to remove partially sequenced, misannotated, and taxonomically overrepresented homologs. BLASTp hits included protein sequences from homologous Mo-, V-, and Fe-nitrogenase isozymes [11]. H-, D, and K-subunit sequences from these isozymes were individually aligned by MAFFT v7.450 [52] and concatenated along with outgroup dark-operative protochlorophyllide oxidoreductase sequences (Bch/ChlLNB). The final dataset included 385 nitrogenase sequences and 385 outgroup sequences. For sequences used to construct Anc1A and Anc2 (internal nodes #960 and #929, respectively), tree reconstruction (using a trimmed alignment generated by trimAl v1.2 [53]) and ancestral sequence inference (using the initial untrimmed alignment) were both performed by RAxML v8.2.10 [54] with the LG+G+F evolutionary model (model testing performed by the ModelFinder [55] in the IQ-TREE v.1.6.12 package [56]).

Due to concerns that RAxML v.8.2 does not implement full, marginal ancestral sequence reconstruction as described by Yang et al. [57], we performed a second phylogenetic analysis as follows. The extant sequence dataset described above was realigned by MAFFT (untrimmed) and tree reconstruction was again performed by RAxML. Ancestral sequence reconstruction was instead performed by PAML v4.9j [58] using the LG+G+F model. From this second reconstruction, Anc1B (internal node #1312), equivalent to Anc1A, was selected for experimental analysis. Anc1B and Anc1A have identical sets of descendent homologs, and their NifD proteins are 95% identical (**Fig. 2*B***).

Only the ancestral sequences inferred with the most probable residue at each protein site were considered for this study (mean posterior probabilities of targeted nitrogenase subunits range from .95 to .99; see **Supplementary File 1a**). All ancestral sequences were reconstructed from well-supported clades (SH-like aLRT = 99-100 [59]).

### Ancestral nitrogenase structural modeling and sequence analysis

Structural homology models of ancestral sequences were generated by MODELLER v10.2 [60] using PDB 1M34 as a template for all nitrogenase protein subunits and visualized by ChimeraX v1.3 [61].

Extant and ancestral protein sequence space was visualized by machine-learning embeddings, where each protein embedding represents protein features in a fixed-size, multidimensional vector space. The analysis was conducted on concatenated (HDK) nitrogenase protein sequences in our phylogenetic dataset. The embeddings were obtained using the pre-trained language model ESM2 [62, 63], a transformer architecture trained to reproduce correlations at the sequence level in a dataset containing hundreds of millions of protein sequences. Layer 33 of this transformer was used, as recommended by the authors. The resulting 1,024 dimensions were reduced by UMAP [64] for visualization in a two-dimensional space.

Protein site-wise conservation analysis was performed using the Consurf server [38]. An input alignment containing only extant, Group I Mo-nitrogenases was submitted for analysis under default parameters. Conserved sites were defined by a Consurf conservation score >7.

### *A. vinelandii* strain engineering

Nucleotide sequences of targeted ancestral nitrogenase proteins were codon-optimized for *A. vinelandii* by a semi-randomized strategy that maximized ancestral nucleotide sequence identity to WT genes. Ancestral and WT protein sequences were compared using the alignment output of ancestral sequence reconstruction (RAxML or PAML). For sites where the ancestral and WT residues were identical, the WT codon was assigned. At sites where the residues were different, the codon was assigned randomly, weighted by *A. vinelandii* codon frequencies (Codon Usage Database, https://www.kazusa.or.jp/codon/). Nucleotide sequences were synthesized into XbaI/KpnI sites of pUC19 vectors (unable to replicate in *A. vinelandii*) (Twist Bioscience; GenScript). Inserts were designed with 400-base-pair flanking regions for homology-directed recombination at the relevant *A. vinelandii nif* locus. An “ASWSHPQFEK” Strep-II-tag was included at the N-terminus of each synthetic *nifD* gene for downstream NifD immunodetection and NifDK affinity purification. See **Supplementary File 1c** for a list of strains and plasmids used in this study.

Engineering of *A. vinelandii* strains used established methods, following Dos Santos [65]. *A. vinelandii* WT (“DJ”), DJ2278 (Δ*nifD::*KanR), DJ2102 (Strep-II-tagged WT NifD), and DJ884 (NifH-overexpression mutant) strains were generously provided by Dennis Dean (Virginia Tech) (**Supplementary File 1c**). Strains Anc1A and Anc2 were constructed from the DJ2278 parent strain via transformation with plasmids pAG13 and pAG19, respectively (**Supplementary File 1c**). For construction of strain Anc2, we first generated a *ΔnifHDK* strain, AK022, by transforming the DJ strain with pAG25. Genetic competency was induced by subculturing relevant parent strains in Mo- and Fe-free Burk’s medium (see below). Competent cells were transformed with at least 1 μg of donor plasmid. Transformants were screened on solid Burk’s medium for rescue of the diazotrophic phenotype (“Nif+”) and loss of kanamycin resistance, followed by Sanger sequencing of the PCR-amplified *nifHDK* cluster (see **Supplementary File 1d** for a list of primers). Transformants were passaged at least 3 times to ensure phenotypic stability prior to storage at −80 °C in phosphate buffer containing 7% DMSO.

### *A. vinelandii* culturing and growth analysis

*A. vinelandii* strains were grown diazotrophically in nitrogen-free Burk’s medium (containing 1 μM Na_2_MoO_4_) at 30 °C and agitated at 300 rpm. To induce genetic competency for transformation experiments, Mo and Fe salts were excluded. For transformant screening, kanamycin antibiotic was added to solid Burk’s medium at a final concentration of 0.6 μg/mL. 50 mL seed cultures for growth rate and acetylene reduction rate quantification were grown non-diazotrophically in flasks with Burk’s medium containing 13 mM ammonium acetate.

For growth rate quantification, seed cultures were inoculated into 100 mL nitrogen-free Burk’s medium to an optical density of ∼0.01 at 600 nm (OD600), after Carruthers et al. [66], and monitored for 72 h. Growth parameters were modeled using the R package Growthcurver [67].

### Microbial acetylene reduction assays

*A. vinelandii* seed cultures representing independent biological replicates were prepared as described above and used to inoculate 100 mL of nitrogen-free Burk’s medium to an OD600 ≈ 0.01. Cells were grown diazotrophically to an OD600 ≈ 0.5, at which point a rubber septum cap was affixed to the mouth of each flask. 25 mL of headspace was removed and replaced by injecting an equivalent volume of acetylene gas. The cultures were subsequently shaken at 30 °C and agitated at 300 rpm. Headspace samples were taken after 15, 30, 45, and 60 mins of incubation for ethylene quantification by a Nexis GC-2030 gas chromatograph (Shimadzu). After the 60 min incubation period, cells were pelleted at 4,700 rpm for 10 mins, washed once with 4 mL of phosphate buffer, and pelleted once more under the same conditions prior to storage at - 80 °C. Total protein was quantified using the Quick Start™ Bradford Protein Assay kit (Bio-Rad) according to manufacturer instructions and a CLARIOstar Plus plate reader (BMG Labtech). Acetylene reduction rates for each replicate were normalized to total protein.

### Nitrogenase expression analysis

Strep-II-tagged NifD protein quantification was performed on all ancestral strains (Anc1A, Anc1B, Anc2) and DJ2102 (harboring Strep-II-tagged WT NifD). Diazotrophic *A. vinelandii* cultures (100 mL) representing three independent biological replicates were prepared as described above and harvested at an OD600 ≈ 1. Cell pellets were resuspended in TE lysis buffer (10 mM Tris, 1 mM EDTA, 1 mg/mL lysozyme) and heated at 95 °C for 10 min. Cell lysates were centrifuged at 5,000 rpm for 15 min. Total protein in the resulting supernatant was quantified using the Pierce™ BCA Protein Assay kit (ThermoFisher) following manufacturer instructions. Normalized protein samples were diluted in 2 × Laemmli buffer at a 1:1 (v/v) ratio prior to SDS-PAGE analysis. Proteins were transferred to nitrocellulose membranes (ThermoFisher), stained with Revert 700 Total Protein Stain (LI-COR), and imaged on a Odyssey Fc Imager (LI-COR). Membranes were then destained with Revert Destaining Solution (LI-COR) and blocked with 5% non-fat milk in PBS solution (137 mM NaCl, 2.7 mM KCl, 10 mM Na_2_HPO_4_, 1.8 mM KH_2_PO_4_.) for 1 h at room temperature. Membranes were rinsed once with PBS-T (PBS with 0.01% Tween-20) and incubated with primary Strep-tag II antibody (Strep-MAB-Classic, IBA Lifesciences, Cat# 2-1507-001, RRID: AB_513133; 1:5000 in 0.2% BSA) for 2 h at room temperature. Membranes were then incubated in LI-COR blocking buffer containing 1:15000 IRDye 680RD Goat anti-Mouse (LI-COR) for 2 h at room temperature and subsequently imaged with an Odyssey Fc Imager (LI-COR). Densitometry analysis was performed with ImageJ [68], with Strep-II-tagged NifD signal intensity normalized to that of the total protein stain.

### Nitrogenase expression, purification, and biochemical characterization

Ancestral nitrogenase NifDK proteins were expressed from relevant *A. vinelandii* strains (Anc1A, Anc1B, Anc2) and purified according to previously published methods [69] with the following modifications: cells were grown diazotrophically in nitrogen-free Burk’s medium and no derepression step to a sufficient OD600 (∼1.8) before harvesting. WT NifH was expressed in *A. vinelandii* strain DJ884 and purified by previously published methods [70]. Protein purity was assessed at ≥ 95% by SDS-PAGE gel with Coomassie blue staining (**Fig. 4-Figure Supplement 1**).

Assays were performed in 9.4 mL vials with a MgATP regeneration buffer (6.7 mM MgCl2, 30 mM phosphocreatine, 5 mM ATP, 0.2 mg/mL creatine phosphokinase, 1.2 mg/mL BSA) and 10 mM sodium dithionite in 100 mM MOPS buffer at pH 7.0. Reaction vials were made anaerobic and relevant gases (N_2_, C_2_H_2_, H_2_) were added to appropriate concentrations with the headspace balanced by argon. NifDK proteins (∼240 kDa) were added to 0.42 µM, the vial vented to atmospheric pressure, and the reaction initiated by addition of NifH (∼60 kDa) protein to 8.4 µM. Reactions were run, shaking, at 30° C for 8 minutes and stopped by the addition of 500 µL of 400 mM EDTA pH 8.0. NH_3_ was quantified using a fluorescence protocol [71] with the following modifications: an aliquot of the sample was added to a solution containing 200 mM potassium phosphate pH 7.3, 20 mM o-phthalaldehyde, and 3.5 mM 2-mercaptoethanol and incubated for 30 minutes in the dark. Fluorescence was measured at λ_excitation_ of 410 nm and λ_emission_ of 472 nm and NH_3_ was quantified using a standard generated with NH_4_Cl. H_2_ and C_2_H_4_ were quantified by gas chromatography by a thermal conductivity detector (GC-TCD) and gas chromatography with flame ionization detector (GC-FID) respectively, according to published methods [72, 73].

### Statistical analyses

Experimental data was statistically analyzed by one-way ANOVA with post-hoc Tukey HSD test.

## MATERIALS AVAILABILITY

Physical materials including bacterial strains and plasmids are available to the scientific community upon request.

## DATA AND CODE AVAILABILITY

Phylogenetic data, including sequence alignments and phylogenetic trees, and the script for ancestral gene codon-optimization are publicly available at https://github.com/kacarlab/garcia_nif2023. All other data are included as source data and supplementary files.

## ACKNOWLEDGEMENTS

We thank Dennis Dean and Valerie Cash for providing *A. vinelandii* strains DJ, DJ2278, DJ2102, and DJ884 and for guidance in genomic manipulations; Jean-Michel Ané, April MacIntyre, and Junko Maeda for guidance and instrumentation support in performing the *in vivo* acetylene reduction assays; Bruno Cuevas for assistance with visualizing nitrogenase sequence space; and the members of the Metal Selection and Utilization Across Eons (MUSE) Consortium for helpful suggestions and discussions. This research was supported by the National Aeronautics and Space Administration (NASA) Interdisciplinary Consortium for Astrobiology Research: Metal Utilization and Selection Across Eons, MUSE (19-ICAR19_2-0007), the University of Wisconsin-Madison College of Agricultural and Life Sciences, the NASA Postdoctoral Program (A.K.G.), NASA Arizona Space Grant (B.M.C.), and the NASA Early Career Faculty Award (B.K.).

## SUPPLEMENTARY FILES & LEGENDS

garcia_elife_2023_suppfile1.pdf: **Supplementary File 1**

**Supplementary File 1a. Sequence characteristics of ancestral nitrogenase subunits**.

**Supplementary File 1b. Host taxa of nitrogenase and outgroup dark-operative protochlorophyllide oxidoreductase homologs included for phylogenetic analysis**.

**Supplementary File 1c. Strains and plasmids used in this study**.

**Supplementary File 1d. Primers used in this study**.

## TABLES

n/a

---

**Figure 2-Figure Supplement 1.**
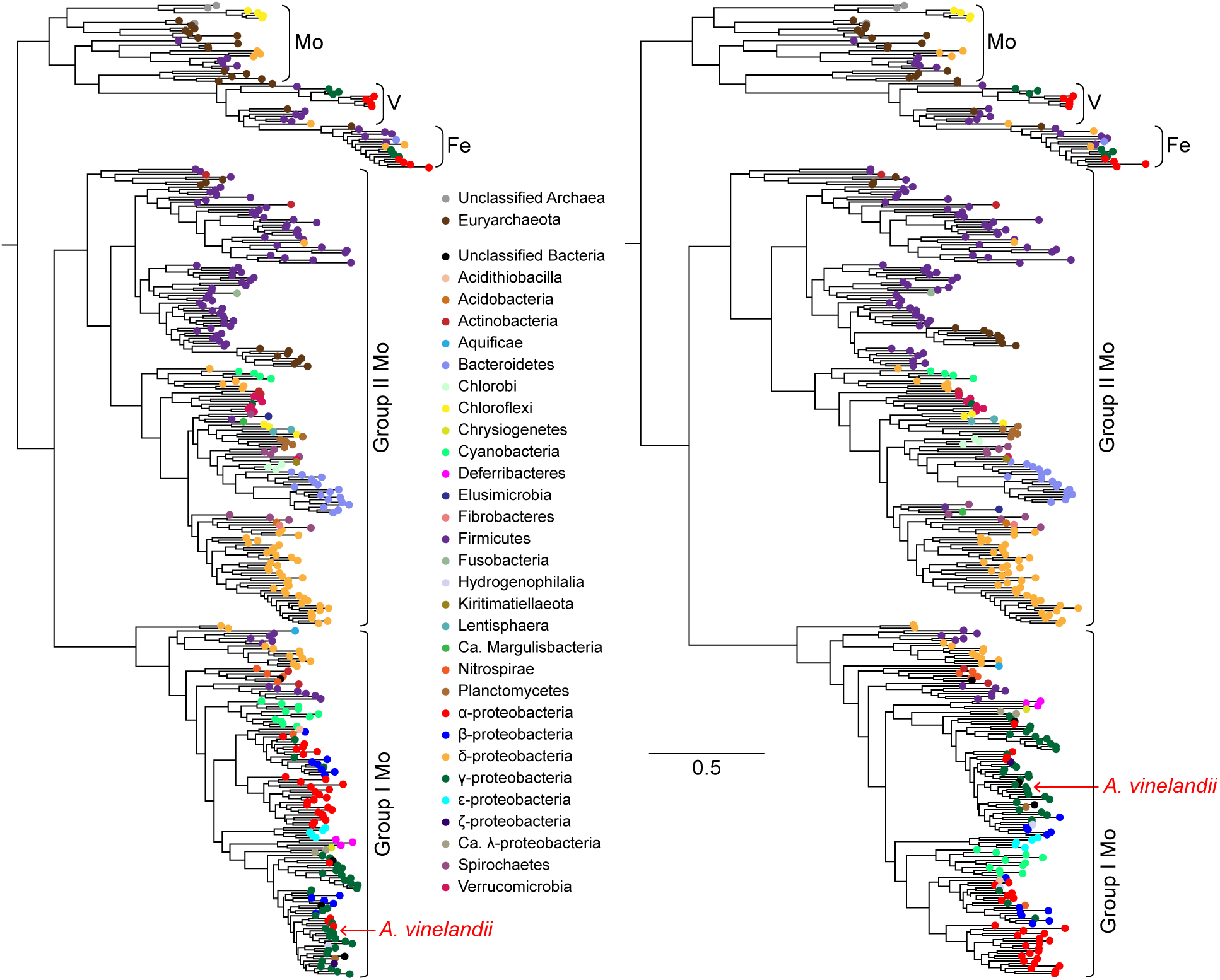
Maximum-likelihood phylogenies built from nitrogenase NifHDK homologs. Anc1A and Anc2 sequences were inferred from the left tree, and the Anc1B sequence was inferred from the right tree (targeting an equivalent node to Anc1A). Both trees were reconstructed from the same extant sequence dataset (see **Materials and methods** for a description of phylogenetic reconstruction and ancestral sequence inference methods). Branch length scale indicates amino acid substitutions per site and applies to both trees.

