## Supplemental File for "Nitrogenase resurrection and the evolution of a singular enzymatic mechanism"

### SUPPLEMENTARY INFORMATION

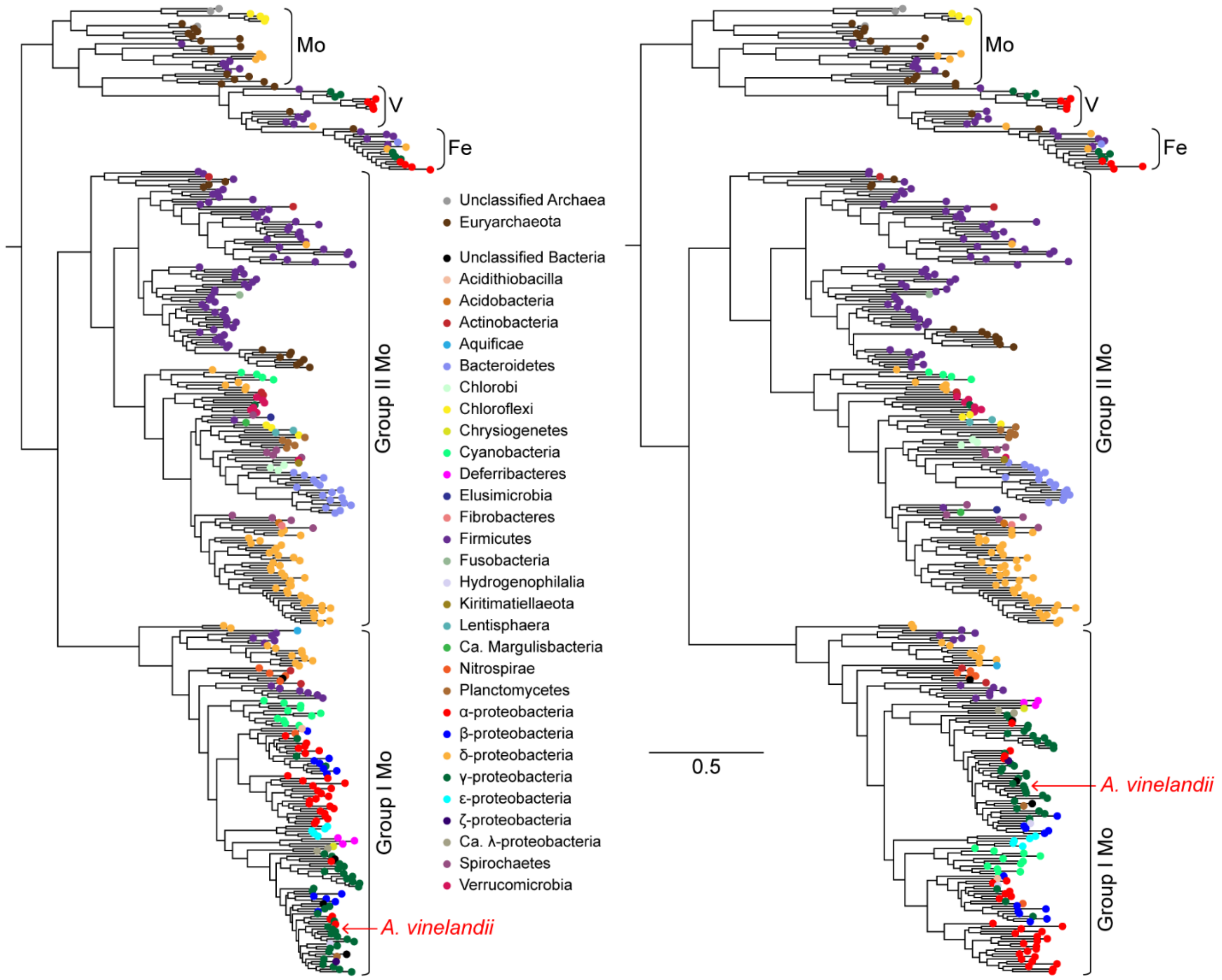


**Figure S1. Maximum-likelihood phylogenies built from nitrogenase NifHDK homologs.** Anc1A and Anc2 sequences were inferred from the left tree, and the Anc1B sequence was inferred from the right tree (targeting an equivalent node to Anc1A). Both trees were reconstructed from the same extant sequence dataset (see **Materials and methods** for a description of phylogenetic reconstruction and ancestral sequence inference methods). Branch length scale indicates amino acid substitutions per site and applies to both trees.

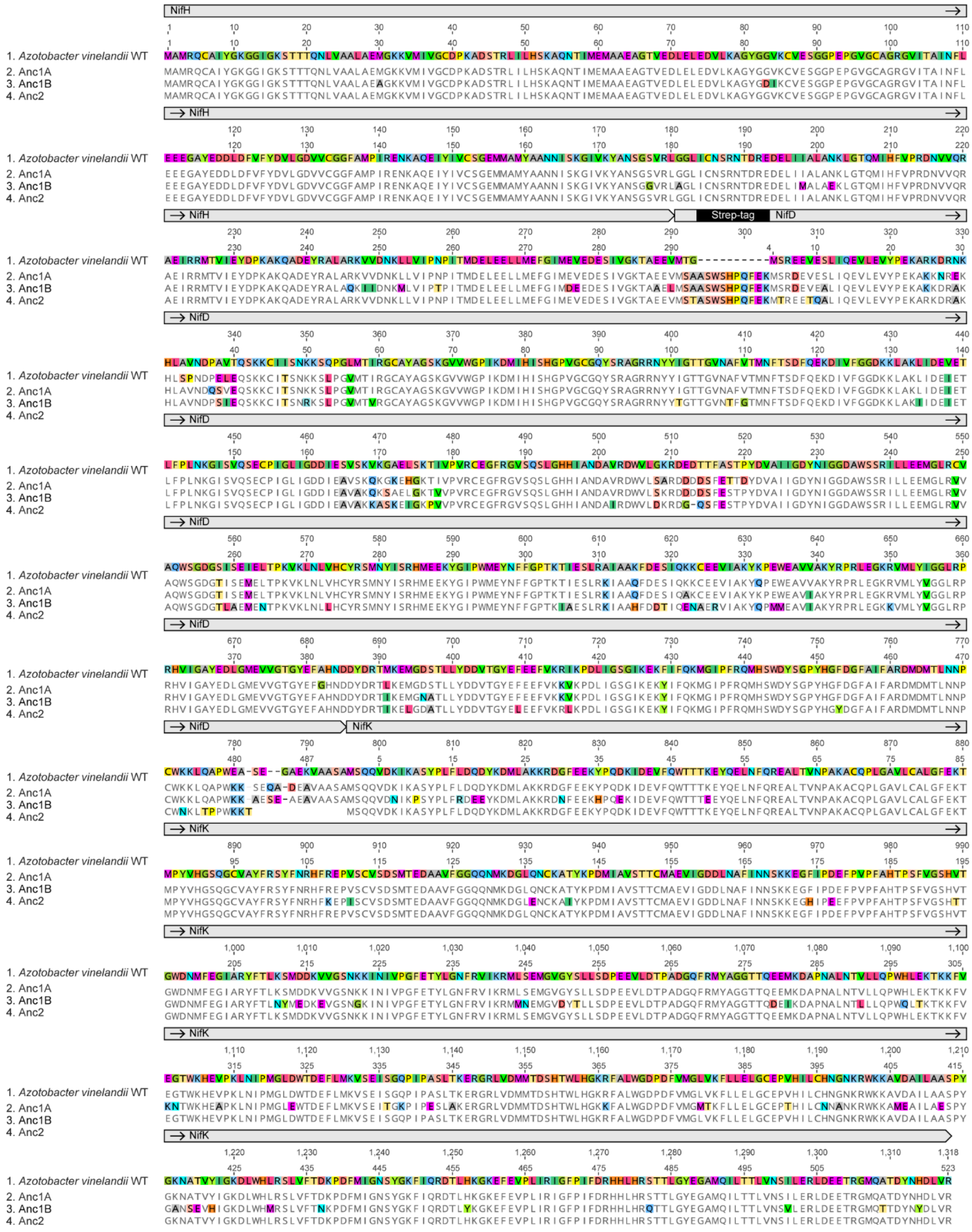


**Figure S2. Alignment of WT and ancestral NifHDK nitrogenase proteins.**

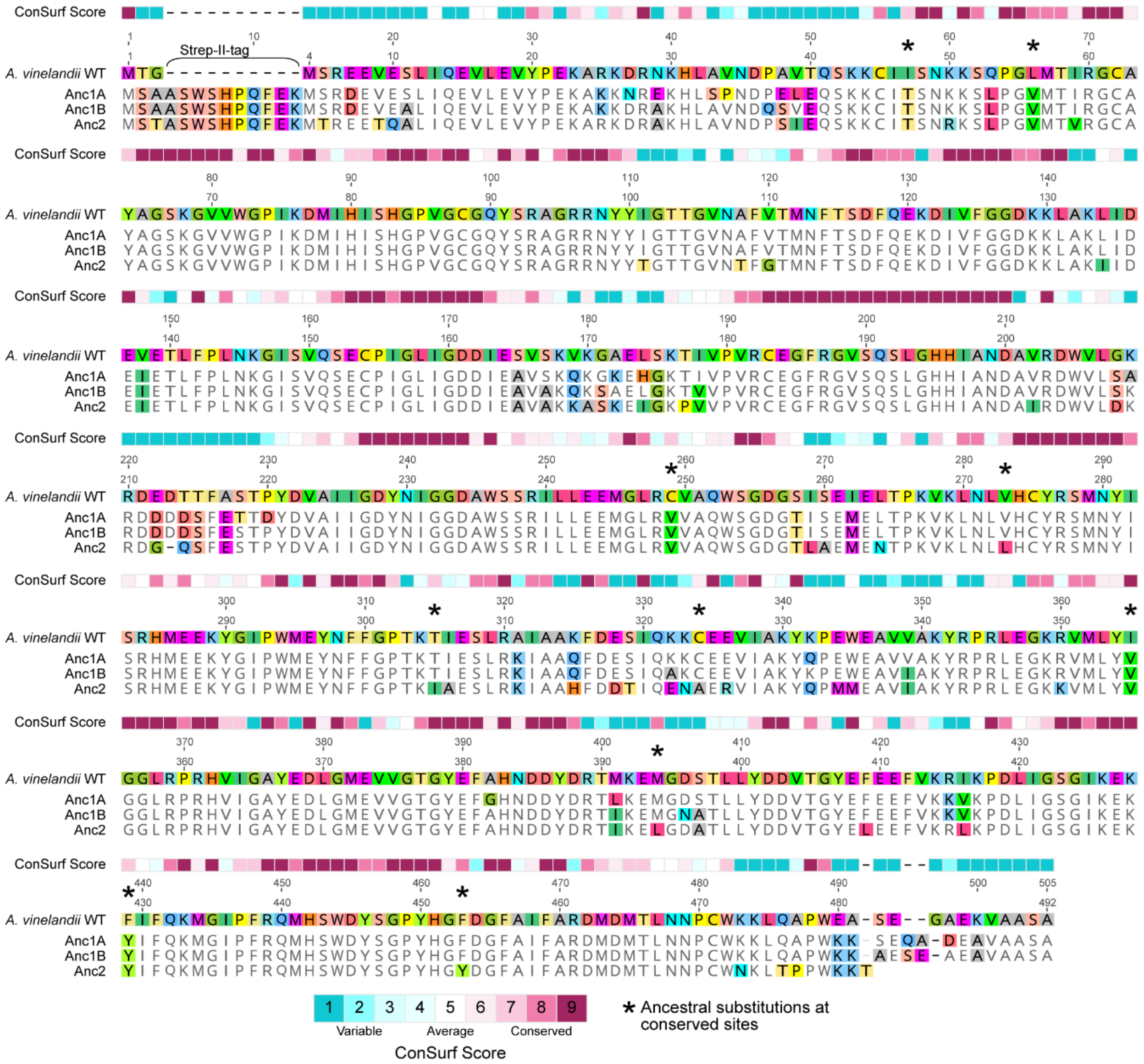

**Figure S3. WT and ancestral NifD alignment with sitewise ConSurf conservation scores (see Materials and Methods).** Conserved sites defined by a ConSurf conservation score >7.

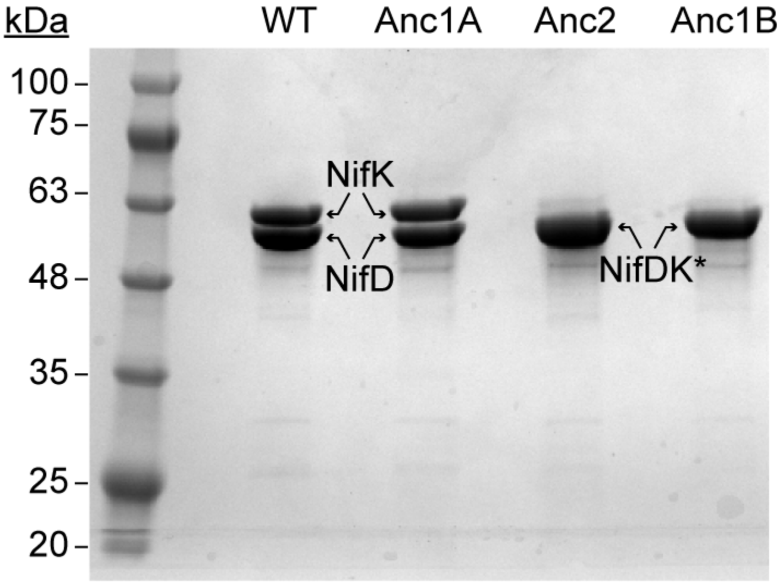


**Figure S4. SDS-PAGE of purified WT and ancestral NifDK proteins. ***NifD and NifK were not separately resolved for Anc2 and Anc1B. Qualitative assessment of band density suggests both subunits migrate together. The presence of both NifD and NifK is inferred based on observed $N_2$ reduction activity of these fractions together with WT NifH (see **Fig 4**).

**Figure S4–Source Data 1. Zip archive of SDS-PAGE image data, containing labeled and unlabeled image files.**

**Table S1. Sequence characteristics of ancestral nitrogenase subunits.**

| Subunit | % protein identity to WT | Mean site posterior probability | % sites with plausible alternate ancestral states | Substitutions at relatively conserved sites† |
|---|---|---|---|---|
| $NifH^{Anc1B}$ | 94% | 0.99 | 1% | G181A |
| $NifD^{Anc1A,Anc1B}$ | 89% | 0.97 | 3% | I47T, L56V**, C249V**, I355V*, F429Y** |
| $NifK^{Anc1B}$ | 89% | 0.98 | 2% | R108K**, H396N** |
| $NifD^{Anc2}$ | 83% | 0.95 | 2% | I47T, L56V**, C249V**, V273L, T305I, C324A, I355V*, M394L, F429Y**, F453Y |

*Active-site

**NifD:NifK interface

†Defined by a Consurf score > 7; see Materials and Methods

**Table S2. Host taxa of nitrogenase and outgroup dark-operative protochlorophyllide oxidoreductase homologs included for phylogenetic analysis.**

| Type | Organism |
|---|---|
| Nif | Acetobacter peroxydans |
| Nif | Acetobacterium woodii |
| Nif | Acidiferrobacter sp. SPIII_3 |
| Nif | Acidihalobacter prosperus |
| Nif | Acidithiobacillus ferridurans |
| Nif | Actibacterium ureilyticum |
| Nif | Actinobacteria bacterium HGW-Actinobacteria-2 |
| Nif | Actinobacteria bacterium HGW-Actinobacteria-10 |
| Nif | Acuticoccus sp. PTG4-2 |
| Nif | Afifella pfennigii |
| Nif | Agaribacterium haliotis |
| Nif | Aliagarivorans marinus |
| Nif | Alkalibacter saccharofermentans |
| Nif | Alkaliflexus imshenetskii |
| Nif | Alkaliphilus metalliredigens |
| Nif | Alphaproteobacteria bacterium |
| Nif | Alteromonadaceae bacterium Bs31 |
| Nif | Alteromonadales bacterium BS08 |
| Nif | Ammonifex sp. |
| Nif | Amphritea atlantica |
| Nif | Anaeroarcus burkinensis |
| Nif | Anaerobacillus alkalidiazotrophicus |
| Nif | Anaerobacterium chartisolvens |
| Nif | Anaerocolumna jejuensis |
| Nif | Anaeromyxobacter sp. Fw109-5 |
| Nif | Anaerospora hongkongensis |
| Nif | Anaerosporobacter mobilis |
| Nif | Anaerosporomusa subterranea |
| Nif | Anaerotignum neopropionicum |

| | |
|---|---|
| Nif | Aneurinibacillus terranovensis |
| Nif | ANME-2 cluster archaeon |
| Nif | archaeon BMS3Abin16 |
| Nif | archaeon BMS3Bbin15 |
| Nif | Arcticibacter svalbardensis |
| Nif | Azoarcus sp. CC-YHH838 |
| Nif | Azotobacter vinelandii |
| Nif | Bacillus caseinilyticus |
| Nif | bacterium BMS3Abin10 |
| Nif | bacterium BMS3Abin12 |
| Nif | bacterium BMS3Abin13 |
| Nif | bacterium BMS3Bbin07 |
| Nif | Bacteroidales bacterium |
| Nif | Bacteroidales bacterium Barb6XT |
| Nif | Bacteroides graminisolvens |
| Nif | Bacteroides luti |
| Nif | Beggiatoa alba |
| Nif | Betaproteobacteria bacterium HGW-Betaproteobacteria-11 |
| Nif | Biostraticola sp. BGMRC 2031 |
| Nif | Blastochloris sp. GI |
| Nif | Blastopirellula marina |
| Nif | Bradyrhizobium oligotrophicum |
| Nif | Caldicellulosiruptor saccharolyticus |
| Nif | Calditerrivibrio nitroreducens |
| Nif | Candidatus Accumulibacter aalborgensis |
| Nif | Candidatus Achromatium palustre |
| Nif | Candidatus Acididesulfobacter diazotrophicus |
| Nif | Candidatus Acidulodesulfobacterium acidiphilum |
| Nif | Candidatus Argoarchaeum ethanivorans |
| Nif | Candidatus Azobacteroides pseudotrichonymphae |
| Nif | Candidatus Competibacteraceae bacterium |
| Nif | Candidatus Dadabacteria bacterium |
| Nif | Candidatus Electronema sp. GS |

| | |
|---|---|
| Nif | Candidatus Lambdaproteobacteria bacterium RIFOXYC1_FULL_56_13 |
| Nif | Candidatus Lambdaproteobacteria bacterium RIFOXYD2_FULL_50_16 |
| Nif | Candidatus Magnetoovum chiemensis |
| Nif | Candidatus Margulisbacteria bacterium GWD2_39_127 |
| Nif | Candidatus Marispirochaeta associata |
| Nif | Candidatus Methanolliviera hydrocarbonicum |
| Nif | Candidatus Methanolliviera sp. GoM_asphalt |
| Nif | Candidatus Methanoperedens nitroreducens |
| Nif | Candidatus Methanoperedens sp. BLZ2 |
| Nif | Candidatus Thiosymbion oneisti |
| Nif | Candidatus Viridilinea mediisalina |
| Nif | Carboxydocella sp. ULO1 |
| Nif | Celerinatantimonas diazotrophica |
| Nif | Cellulosilyticum lentocellum |
| Nif | Chlorobium chlorochromatii |
| Nif | Chloroflexales bacterium ZM16-3 |
| Nif | Chloroflexi bacterium |
| Nif | Chloroherpeton thalassium |
| Nif | Chromatiaceae bacterium 2141T.STBD.0c.01a |
| Nif | Chrysiogenes arsenatis |
| Nif | Clostridiales bacterium 43-6 |
| Nif | Clostridiales bacterium DRI-13 |
| Nif | [Clostridium] algidixylanolyticum |
| Nif | [Clostridium] fimetarium |
| Nif | Clostridium pasteurianum |
| Nif | [Clostridium] populeti |
| Nif | Cohaesibacter haloalkalitolerans |
| Nif | Coleofasciculus chthonoplastes |
| Nif | Coraliomargarita akajimensis |
| Nif | Cupriavidus sp. amp6 |
| Nif | cyanobacterium endosymbiont of Epithemia turgida |

| | |
|---|---|
| Nif | Cytophaga xylanolytica |
| Nif | Defluviimonas alba |
| Nif | Defluviitalea phaphyphila |
| Nif | Dehalobacter |
| Nif | Dehalococcoides mccartyi |
| Nif | Dehalogenimonas sp. WBC-2 |
| Nif | delta proteobacterium NaphS2 |
| Nif | Deltaproteobacteria bacterium |
| Nif | Deltaproteobacteria bacterium HGW-Deltaproteobacteria-4 |
| Nif | Deltaproteobacteria bacterium HGW-Deltaproteobacteria-8 |
| Nif | Dendrosporobacter quercicolus |
| Nif | Derxia gummosa |
| Nif | Desertifilum sp. IPPAS B-1220 |
| Nif | Desulfacinum infernum |
| Nif | Desulfallas arcticus |
| Nif | Desulfamplus magnetovallimortis |
| Nif | Desulfatibacillum aliphaticivorans |
| Nif | Desulfatitalea sp. BRH_c12 |
| Nif | Desulfobacca acetoxidans |
| Nif | Desulfobacteraceae bacterium |
| Nif | Desulfobacteraceae bacterium Eth-SRB1 |
| Nif | Desulfobacteraceae bacterium Eth-SRB2 |
| Nif | Desulfobacterales bacterium C00003106 |
| Nif | Desulfobacterium autotrophicum |
| Nif | Desulfobotulus alkaliphilus |
| Nif | Desulfobulbaceae bacterium |
| Nif | Desulfobulbaceae bacterium A2 |
| Nif | Desulfobulbaceae bacterium DB1 |
| Nif | Desulfobulbus elongatus |
| Nif | Desulfocarbo indianensis |
| Nif | Desulfocucumis palustris |
| Nif | Desulfocurvibacter africanus |
| Nif | Desulfofarcimen acetoxidans |

| | |
|---|---|
| Nif | Desulfofustis glycolicus |
| Nif | Desulfoglaeba alkanexedens |
| Nif | Desulfohalovibrio alkalitolerans |
| Nif | Desulfoluna spongiiphila |
| Nif | Desulfomicrobium apsheronum |
| Nif | Desulfomonile tiedjei |
| Nif | Desulfonatronospira thiodismutans |
| Nif | Desulfonatronovibrio hydrogenovorans |
| Nif | Desulfonatronum lacustre |
| Nif | Desulfopila aestuarii |
| Nif | Desulforegula conservatrix |
| Nif | Desulforhopalus singaporensis |
| Nif | Desulfosarcina cetonica |
| Nif | Desulfospira joergensenii |
| Nif | Desulfosporosinus fructosivorans |
| Nif | Desulfotalea sp. |
| Nif | Desulfotomaculum ferrireducens |
| Nif | Desulfovibrio desulfuricans |
| Nif | Desulfuribacillus alkaliarsenatis |
| Nif | Desulfurispora thermophila |
| Nif | Desulfurobacterium atlanticum |
| Nif | Desulfuromonadales bacterium GWC2_61_20 |
| Nif | Desulfuromonas acetexigens |
| Nif | Dethiobacter alkaliphilus |
| Nif | Dethiosulfatibacter aminovorans |
| Nif | Dissulfuribacter thermophilus |
| Nif | Draconibacterium sediminis |
| Nif | Dysgonomonas capnocytophagoides |
| Nif | Ectothiorhodospira haloalkaliphila |
| Nif | Ectothiorhodospiraceae bacterium |
| Nif | Elusimicrobia bacterium RIFOXYA2_FULL_50_26 |
| Nif | Ethanoligenens harbinense |
| Nif | Euhalothece natronophila |

| | |
|---|---|
| Nif | Fibrobacteres bacterium |
| Nif | filamentous cyanobacterium ESFC-1 |
| Nif | Firmicutes bacterium HGW-Firmicutes-1 |
| Nif | Firmicutes bacterium HGW-Firmicutes-8 |
| Nif | Fischerella sp. PCC 9605 |
| Nif | Fontibacillus panacisegetis |
| Nif | Frankia canadensis |
| Nif | Gaiellales bacterium |
| Nif | Gallionellales bacterium GWA2_60_18 |
| Nif | Gammaproteobacteria bacterium 28-57-27 |
| Nif | Geitlerinema sp. PCC 9228 |
| Nif | Geminicoccaceae bacterium |
| Nif | Geminisphaera colitermitum |
| Nif | Geoalkalibacter ferrihydriticus |
| Nif | Geobacter thiogenes |
| Nif | Geosporobacter ferrireducens |
| Nif | Geothermobacter sp. EPR-M |
| Nif | Geovibrio sp. L21-Ace-BES |
| Nif | Gloeocapsa sp. DLM2.Bin57 |
| Nif | Gordonibacter sp. 28C |
| Nif | Gorillibacterium timonense |
| Nif | Gracilibacter sp. BRH_c7a |
| Nif | Halanaerobium salsuginis |
| Nif | Haliea sp. SAOS-164 |
| Nif | Halocella sp. SP3-1 |
| Nif | Halorhodospira halochloris |
| Nif | Hartmannibacter diazotrophicus |
| Nif | Heliobacterium modesticaldum |
| Nif | Herbaspirillum frisingense |
| Nif | Holophaga foetida |
| Nif | Hungateiclostridium cellulolyticum |
| Nif | Hydrogenispora ethanolica |
| Nif | Hydrogenophilales bacterium 16-64-40 |

| | |
|---|---|
| Nif | Hydrogenophilales bacterium 28-61-23 |
| Nif | Ilyobacter polytropus |
| Nif | Kamptonema |
| Nif | Kiritimatiellales bacterium |
| Nif | Klebsiella pneumoniae |
| Nif | Labilibacter aurantiacus |
| Nif | Lachnospiraceae bacterium |
| Nif | Lachnospiraceae bacterium Marseille-P3773 |
| Nif | Lebetimonas sp. JS032 |
| Nif | Lentisphaerae bacterium GWF2_50_93 |
| Nif | Lentisphaerae bacterium GWF2_57_35 |
| Nif | [Leptolyngbya] sp. JSC-1 |
| Nif | Leptospirillum ferriphilum |
| Nif | Lucifera butyrica |
| Nif | Lutibacter agarilyticus |
| Nif | Lyngbya sp. PCC 8106 |
| Nif | magneto-ovoid bacterium MO-1 |
| Nif | Magnetovibrio blakemorei |
| Nif | Malonomonas rubra |
| Nif | Mangrovibacter sp. MFB070 |
| Nif | Mangrovibacterium diazotrophicum |
| Nif | Marinilabilia sp. WTE |
| Nif | Marinobacter sp. ES-1 |
| Nif | Mariprofundus ferrooxydans |
| Nif | Martelella sp. BGMRC2036 |
| Nif | Massilibacillus massiliensis |
| Nif | Megasphaera cerevisiae |
| Nif | Methanobacteriales archaeon HGW-Methanobacteriales-1 |
| Nif | Methanobacterium paludis |
| Nif | Methanobrevibacter curvatus |
| Nif | Methanocaldococcus infernus |
| Nif | Methanocella arvoryzae |
| Nif | Methanococcus maripaludis |

| | |
|---|---|
| Nif | Methanoculleus taiwanensis |
| Nif | Methanolacinia paynteri |
| Nif | Methanolobus psychrotolerans |
| Nif | Methanomassiliicoccus luminyensis |
| Nif | Methanomicrobiales archaeon HGW-Methanomicrobiales-1 |
| Nif | Methanophagales archaeon |
| Nif | Methanoregula boonei |
| Nif | Methanosarcina acetivorans C2A |
| Nif | Methanosphaerula palustris |
| Nif | Methanospirillum lacunae |
| Nif | Methanothermobacter thermautotrophicus |
| Nif | Methanothermococcus okinawensis |
| Nif | Methanothrix soehngenii |
| Nif | Methanotorris igneus |
| Nif | Methyloferula stellata |
| Nif | Methyloglobulus morosus |
| Nif | Methylomonas koyamae |
| Nif | Methyloprofundus sedimenti |
| Nif | Moorella thermoacetica |
| Nif | Natranaerovirga hydrolytica |
| Nif | Natronoflexus pectinivorans |
| Nif | Neorhizobium galegae |
| Nif | Nitrospirae bacterium |
| Nif | Nitrospirae bacterium GWC2_57_13 |
| Nif | Oceanospirillaceae bacterium |
| Nif | Opitutaceae bacterium EW11 |
| Nif | Orenia marismortui |
| Nif | Orenia metallireducens |
| Nif | Oscillochloris trichoides |
| Nif | Oxalobacter sp. |
| Nif | Oxobacter pfennigii |
| Nif | Paludibacter propionicigenes |
| Nif | Pantoea cypripedii |

| | |
|---|---|
| Nif | Parasporobacterium paucivorans |
| Nif | Pectinatus cerevisiiphilus |
| Nif | Pelobacter acetylenicus |
| Nif | Pelomonas saccharophila |
| Nif | Pelotomaculum sp. FP |
| Nif | Peptococcaceae bacterium BICA1-7 |
| Nif | Peptococcaceae bacterium BRH_c4a |
| Nif | Peptococcaceae bacterium CEB3 |
| Nif | Peptococcaceae bacterium SCADC1_2_3 |
| Nif | Petrocella atlantisensis |
| Nif | Petroclostridium xylanilyticum |
| Nif | Phaeospirillum fulvum |
| Nif | Phormidium sp. HE10JO |
| Nif | Phycisphaeraceae bacterium |
| Nif | Planctomycetaceae bacterium |
| Nif | Planctomycetes bacterium |
| Nif | Planctomycetes bacterium Q31b |
| Nif | Planctomycetes bacterium V7 |
| Nif | Planktothrix tepida |
| Nif | Pleomorphomonas carboxyditropha |
| Nif | Polaromonas naphthalenivorans |
| Nif | Prevotella oryzae |
| Nif | Propionibacterium cyclohexanicum |
| Nif | Propionispira arboris |
| Nif | Propionivibrio dicarboxylicus |
| Nif | Prosthecochloris sp. GSB1 |
| Nif | Proteobacteria bacterium CG1_02_64_396 |
| Nif | Pseudobacteroides cellulosolvens |
| Nif | Pseudoclostridium thermosuccinogenes |
| Nif | Pseudodesulfovibrio hydrargyri |
| Nif | Rhizobiales bacterium |
| Nif | Rhodobacteraceae bacterium |
| Nif | Rhodoblastus acidophilus |

| | |
|---|---|
| Nif | Rhodocyclaceae bacterium |
| Nif | Rhodopila globiformis |
| Nif | Rhodopseudomonas palustris |
| Nif | Rhodospirillaceae bacterium BRH_c57 |
| Nif | Rhodothalassium salexigens |
| Nif | Rhodovibrio salinarum |
| Nif | Robinsoniella sp. MCWD5 |
| Nif | Roseiflexus castenholzii |
| Nif | Roseofilum reptotaenium AO1-A |
| Nif | Roseospirillum parvum |
| Nif | Ruminiclostridium hungatei |
| Nif | Sediminispirochaeta smaragdinae |
| Nif | Seleniivibrio woodruffii |
| Nif | Serpentinicella alkaliphila |
| Nif | Serratia sp. ATCC 39006 |
| Nif | Skermanella aerolata |
| Nif | Smithella sp. SC_K08D17 |
| Nif | Sodalis sp. 159R |
| Nif | Solimonas aquatica |
| Nif | Spirochaeta cellobiosiphila |
| Nif | Spirochaeta thermophila |
| Nif | Spirochaetaceae bacterium |
| Nif | Spirochaetes bacterium GWB1_27_13 |
| Nif | Spirochaetes bacterium GWB1_36_13 |
| Nif | Spirochaetes bacterium GWE1_32_154 |
| Nif | Sporobacter termitidis |
| Nif | Sporolactobacillus terrae |
| Nif | Sporomusa termitida |
| Nif | Sulfuricurvum kujiense |
| Nif | Sulfurimonas sp. RIFOXYD12_FULL_33_39 |
| Nif | Sulfurivermis fontis |
| Nif | Sulfurovum sp. FS08-3 |
| Nif | Syntrophobacter fumaroxidans |

| | |
|---|---|
| Nif | Syntrophomonas zehnderi |
| Nif | Syntrophorhabdaceae bacterium PtaU1.Bin034 |
| Nif | Syntrophothermus lipocalidus |
| Nif | Syntrophus gentianae |
| Nif | Telmatospirillum siberiense |
| Nif | Terrimicrobium sacchariphilum |
| Nif | Thermacetogenium phaeum |
| Nif | Thermicanus aegyptius |
| Nif | Thermoanaerobacterium thermosaccharolyticum |
| Nif | Thermodesulfitimonas autotrophica |
| Nif | Thermodesulforhabdus norvegica |
| Nif | Thermodesulfovibrio aggregans |
| Nif | Thioflavicoccus mobilis |
| Nif | Thioploca ingrica |
| Nif | Thiorhodococcus drewsii |
| Nif | Tolumonas lignilytica |
| Nif | Treponema primitia |
| Nif | uncultured archaeon |
| Nif | Varunaivibrio sulfuroxidans |
| Nif | Verrucomicrobia bacterium S94 |
| Nif | Verrucomicrobia bacterium Tous-C9LFEB |
| Nif | Verrucomicrobiae bacterium DG1235 |
| Nif | Vibrio aerogenes |
| Nif | Vulcanococcus limneticus |
| Nif | Wolinella succinogenes |
| Nif | Youngiibacter fragilis |
| Nif | Zymomonas mobilis |
| Vnf | Archaeoglobus sp. |
| Vnf | Azomonas agilis |
| Vnf | Azotobacter vinelandii |
| Vnf | Clostridium pasteurianum |
| Vnf | Desulfobacter curvatus |
| Vnf | Ethanoligenens harbinense |

| | |
|---|---|
| Vnf | Lucifera butyrica |
| Vnf | Methanosarcina acetivorans C2A |
| Vnf | Methylocystaceae bacterium |
| Vnf | Methylocystis parvus |
| Vnf | Methylomusa anaerophila |
| Vnf | Paenibacillus durus |
| Vnf | Phaeospirillum fulvum |
| Vnf | Rhodoblastus acidophilus |
| Vnf | Rhodopseudomonas palustris |
| Vnf | Ruminiclostridium hungatei |
| Vnf | Tolumonas lignilytica |
| Anf | Anaerocolumna jejuensis |
| Anf | Azotobacter vinelandii |
| Anf | Clostridium pasteurianum |
| Anf | Desulfosporosinus fructosivorans |
| Anf | Desulfovibrio desulfuricans |
| Anf | Dickeya paradisiaca |
| Anf | Dysgonomonas capnocytophagoides |
| Anf | Geobacter thiogenes |
| Anf | Methanosarcina acetivorans C2A |
| Anf | Pectinatus cerevisiiphilus |
| Anf | Phaeospirillum fulvum |
| Anf | Rahnella sp. AA |
| Anf | Rhodobacter capsulatus |
| Anf | Rhodoblastus acidophilus |
| Anf | Rhodopseudomonas palustris |
| Anf | Sporomusa termitida |
| Anf | Thiorhodococcus drewsii |
| BchChl | Acaryochloris sp. RCC1774 |
| BchChl | Acetobacteraceae bacterium DB1506 |
| BchChl | Acetobacteraceae bacterium KEBCLARHB70R |
| BchChl | Acidibrevibacterium fodinaquatile |
| BchChl | Acidiphilium multivorum |

| | |
|---|---|
| BchChl | Acidocella sp. 20-57-95 |
| BchChl | Acuticoccus kandeliae |
| BchChl | Aestuariivita boseongensis |
| BchChl | Afifella marina |
| BchChl | Agrobacterium albertimagni |
| BchChl | Ahrensia sp. R2A130 |
| BchChl | Albimonas pacifica |
| BchChl | Aliterella atlantica |
| BchChl | Alkalinema sp. CACIAM 70d |
| BchChl | Allochromatium vinosum |
| BchChl | alpha proteobacterium AAP38 |
| BchChl | Alphaproteobacteria bacterium |
| BchChl | Alphaproteobacteria bacterium HGW-Alphaproteobacteria-1 |
| BchChl | Alphaproteobacteria bacterium HGW-Alphaproteobacteria-14 |
| BchChl | Alphaproteobacteria bacterium PA4 |
| BchChl | Alphaproteobacteria bacterium WS11 |
| BchChl | Altererythrobacter ishigakiensis |
| BchChl | Alteromonadaceae bacterium |
| BchChl | Anabaena cylindrica |
| BchChl | Aphanocapsa feldmannii 288cV |
| BchChl | Aphanothece hegewaldii |
| BchChl | Aquabacterium sp. W35 |
| BchChl | Aquidulcibacter paucihalophilus |
| BchChl | Aquincola tertiaricarbonis |
| BchChl | Arthrospira platensis |
| BchChl | Ascidiaceihabitans donghaensis |
| BchChl | Aurantimonas sp. 22II-16-19i |
| BchChl | Azospirillum |
| BchChl | bacterium RmlP026 |
| BchChl | Beijerinckiaceae bacterium RH AL1 |
| BchChl | Belnapia moabensis |
| BchChl | beta proteobacterium AAP51 |
| BchChl | beta proteobacterium AAP99 |

| | |
|---|---|
| BchChl | Betaproteobacteria bacterium HGW-Betaproteobacteria-3 |
| BchChl | Betaproteobacteria bacterium TMED41 |
| BchChl | Betaproteobacteria bacterium TMED82 |
| BchChl | Betaproteobacteria bacterium TMED156 |
| BchChl | Betaproteobacteria bacterium UKL13-2 |
| BchChl | Blastochloris sp. GI |
| BchChl | Blastomonas natatoria |
| BchChl | Bosea sp. AAP35 |
| BchChl | Bradyrhizobiaceae bacterium |
| BchChl | Bradyrhizobium guangzhouense |
| BchChl | Brevundimonas bacteroides |
| BchChl | Burkholderiales bacterium |
| BchChl | Burkholderiales bacterium 28-67-8 |
| BchChl | Burkholderiales bacterium C2 |
| BchChl | Burkholderiales bacterium PBB1 |
| BchChl | Burkholderiales bacterium PBB2 |
| BchChl | Burkholderiales bacterium RIFCSPHIGHO2_12_FULL_69_20 |
| BchChl | Caenispirillum salinarum |
| BchChl | Calothrix brevissima |
| BchChl | Candidatus Atelocyanobacterium thalassa |
| BchChl | Candidatus Chloroploca sp. Khr17 |
| BchChl | Candidatus Oscillochloris fontis |
| BchChl | Candidatus Phycosocius bacilliformis |
| BchChl | Candidatus Synechococcus spongiarum |
| BchChl | Candidatus Thermochlorobacter aerophilum |
| BchChl | Candidatus Thiodictyon syntrophicum |
| BchChl | Candidatus Viridilinea mediisalina |
| BchChl | Caulobacteraceae bacterium PMMR1 |
| BchChl | Cereibacter changlensis |
| BchChl | Chamaesiphon minutus |
| BchChl | Chloracidobacterium thermophilum |
| BchChl | Chlorobaculum limnaeum |
| BchChl | Chlorobium limicola |

| | |
|---|---|
| BchChl | Chloroflexales bacterium ZM16-3 |
| BchChl | Chloroflexus aggregans |
| BchChl | Chlorogloea sp. CCALA 695 |
| BchChl | Chlorogloeopsis fritschii |
| BchChl | Chloroherpeton thalassium |
| BchChl | Chondrocystis sp. NIES-4102 |
| BchChl | Chromatiaceae bacterium |
| BchChl | Chromatium okenii |
| BchChl | Chromatocurvus halotolerans |
| BchChl | Chroococcidiopsis sp. TS-821 |
| BchChl | Chrysosporum ovalisporum |
| BchChl | Citromicrobium sp. WPS32 |
| BchChl | Cognatiyoonia koreensis |
| BchChl | Coleofasciculus chthonoplastes |
| BchChl | Comamonadaceae bacterium |
| BchChl | Comamonadaceae bacterium SYSU G00088 |
| BchChl | Comamonadaceae bacterium YIM 73032 |
| BchChl | Congregibacter litoralis |
| BchChl | Crinalium epipsammum |
| BchChl | Cuspidothrix issatschenkoi |
| BchChl | Cyanobacteria bacterium J007 |
| BchChl | Cyanobacteria bacterium J083 |
| BchChl | Cyanobacteria bacterium UBA12227 |
| BchChl | Cyanobacterium aponinum |
| BchChl | cyanobacterium endosymbiont of Rhopalodia gibberula |
| BchChl | cyanobacterium PCC 7702 |
| BchChl | Cyanobium gracile |
| BchChl | Cyanothece sp. BG0011 |
| BchChl | Cylindrospermum sp. NIES-4074 |
| BchChl | Dactylococcopsis salina |
| BchChl | Dankookia rubra |
| BchChl | Desertifilum sp. IPPAS B-1220 |
| BchChl | Dichotomicrobium thermohalophilum |

| | |
|---|---|
| BchChl | Dinoroseobacter shibae |
| BchChl | Dolichospermum planctonicum |
| BchChl | Dongia mobilis |
| BchChl | Ectothiorhodosinus mongolicus |
| BchChl | Ectothiorhodospira magna |
| BchChl | Elioraea sp. PF-30 |
| BchChl | Erythrobacter litoralis |
| BchChl | Euhalothece natronophila |
| BchChl | filamentous cyanobacterium CCP3 |
| BchChl | filamentous cyanobacterium ESFC-1 |
| BchChl | Filomicrobium sp. |
| BchChl | Fischerella sp. NIES-4106 |
| BchChl | Flavimaricola marinus |
| BchChl | Fortiea contorta |
| BchChl | Fulvimarina manganoxydans |
| BchChl | gamma proteobacterium HIMB55 |
| BchChl | gamma proteobacterium NOR5-3 |
| BchChl | Geitlerinema sp. PCC 7407 |
| BchChl | Geminicoccaceae bacterium |
| BchChl | Geminocystis herdmanii |
| BchChl | Gemmatimonadales bacterium |
| BchChl | Gemmatimonas phototrophica |
| BchChl | Gemmobacter sp. CC-PW-75 |
| BchChl | Gloeobacter kilaueensis |
| BchChl | Gloeocapsa sp. PCC 7428 |
| BchChl | Gloeocapsopsis sp. AAB1 = 1H9 |
| BchChl | Gloeomargarita lithophora |
| BchChl | Gloeothece citriformis |
| BchChl | Halieaceae bacterium |
| BchChl | Halomicronema hongdechloris |
| BchChl | Halorhodospira halochloris |
| BchChl | Halothece sp. PCC 7418 |
| BchChl | Hasllibacter halocynthiae |

| | |
|---|---|
| BchChl | Hassallia byssoidea VB512170 |
| BchChl | Heliobacillus mobilis |
| BchChl | Heliobacterium modesticaldum |
| BchChl | Heliophilum fasciatum |
| BchChl | Histidinibacterium lentulum |
| BchChl | Hoeflea olei |
| BchChl | Hormoscilla sp. GUM007 |
| BchChl | Hwanghaeicola aestuarii |
| BchChl | Hydrococcus rivularis |
| BchChl | Hydrogenophaga flava |
| BchChl | Hyella patelloides |
| BchChl | Hyphomicrobium sp. |
| BchChl | Hyphomicrobium sp. 12-62-95 |
| BchChl | Hyphomicrobium zavarzinii |
| BchChl | Hyphomonadaceae bacterium UKL13-1 |
| BchChl | Ideonella sakaiensis |
| BchChl | Imhoffiella purpurea |
| BchChl | Jannaschia aquimarina |
| BchChl | Kamptonema |
| BchChl | Kandeliimicrobium roseum |
| BchChl | Labrenzia alexandrii |
| BchChl | Leptolyngbya boryana |
| BchChl | [Leptolyngbya] sp. JSC-1 |
| BchChl | Leptothrix mobilis |
| BchChl | Limimaricola pyoseonensis |
| BchChl | Limnohabitans sp. 2KL-17 |
| BchChl | Limnoraphis robusta |
| BchChl | Limnothrix rosea |
| BchChl | Litoreibacter ponti |
| BchChl | Loktanella atrilutea |
| BchChl | Luminiphilus syltensis |
| BchChl | Lutimaribacter saemankumensis |
| BchChl | Lyngbya sp. PCC 8106 |

| | |
|---|---|
| BchChl | Maliponia aquimaris |
| BchChl | Maribius sp. MOLA 401 |
| BchChl | marine gamma proteobacterium HTCC2080 |
| BchChl | Marinibacterium profundimaris |
| BchChl | Marinovum sp. |
| BchChl | Maritimibacter sp. |
| BchChl | Marivita geojedonensis |
| BchChl | Mastigocladopsis repens |
| BchChl | Mastigocoleus testarum |
| BchChl | Merismopedia glauca |
| BchChl | Mesorhizobium loti |
| BchChl | Methylibium sp. NZG |
| BchChl | Methylobacterium brachiatum |
| BchChl | Methylocapsa palsarum |
| BchChl | Methylocella silvestris |
| BchChl | Methylocystis rosea |
| BchChl | Methylorubrum extorquens |
| BchChl | Methyloversatilis sp. RAC08 |
| BchChl | Microcoleus sp. PCC 7113 |
| BchChl | Microcystis aeruginosa |
| BchChl | Mongoliimonas terrestris |
| BchChl | Moorea producens |
| BchChl | Myxococcales bacterium |
| BchChl | Myxosarcina sp. GI1 |
| BchChl | Neosynechococcus sphagnicola |
| BchChl | Nereida ignava |
| BchChl | Nevskia ramosa |
| BchChl | Niveispirillum lacus |
| BchChl | Nodosilinea nodulosa |
| BchChl | Nodularia sp. NIES-3585 |
| BchChl | Nostoc calcicola |
| BchChl | Nostocales cyanobacterium |
| BchChl | Nostocales cyanobacterium HT-58-2 |

| | |
|---|---|
| BchChl | Novosphingobium acidiphilum |
| BchChl | Oceanibaculum indicum |
| BchChl | Oceanicola sp. HL-35 |
| BchChl | Oceanobacter sp. |
| BchChl | Okeania hirsuta |
| BchChl | Oscillatoria acuminata |
| BchChl | Oscillatoria nigro-viridis |
| BchChl | Oscillatoriales cyanobacterium CG2_30_44_21 |
| BchChl | Oscillatoriales cyanobacterium JSC-12 |
| BchChl | Oscillatoriales cyanobacterium USR001 |
| BchChl | Oscillochloris trichoides |
| BchChl | Palleronia abyssalis |
| BchChl | Paracraurococcus ruber |
| BchChl | Pararhizobium sp. BGMRC6574 |
| BchChl | Pararhodospirillum oryzae |
| BchChl | Pelagicola sp. LXJ1103 |
| BchChl | Pelodictyon luteolum |
| BchChl | Pelomonas puraquae |
| BchChl | Phaeobacter sp. 22II1-1F12B |
| BchChl | Phaeospirillum fulvum |
| BchChl | Phormidesmis priestleyi |
| BchChl | Phormidium ambiguum |
| BchChl | Phreatobacter oligotrophus |
| BchChl | Phyllobacteriaceae bacterium |
| BchChl | Phyllobacteriaceae bacterium StC1 |
| BchChl | Phyllobacteriaceae bacterium Z3-1 |
| BchChl | Piscinibacter caeni |
| BchChl | Planktotalea frisia |
| BchChl | Planktothricoides sp. SR001 |
| BchChl | Planktothrix sp. PCC 11201 |
| BchChl | Polymorphobacter sp. DJ1R-1 |
| BchChl | Polynucleobacter acidiphobus |
| BchChl | Ponticoccus marisrubri |

| | |
|---|---|
| BchChl | Pontivivens insulae |
| BchChl | Porphyrobacter colymbi |
| BchChl | Primorskyibacter sp. SS33 |
| BchChl | Prochlorococcus marinus |
| BchChl | Prochlorothrix hollandica |
| BchChl | Propionibacteriaceae bacterium |
| BchChl | Prosthecochloris marina |
| BchChl | Prosthecomicrobium hirschii |
| BchChl | Proteobacteria bacterium SG_bin5 |
| BchChl | Proteobacteria bacterium SG_bin6 |
| BchChl | Proteobacteria bacterium ST_bin13 |
| BchChl | Proteobacteria bacterium ST_bin14 |
| BchChl | Pseudacidovorax sp. RU35E |
| BchChl | Pseudaestuariivita atlantica |
| BchChl | Pseudanabaena biceps |
| BchChl | Pseudohaliea rubra |
| BchChl | Pseudomonas sp. SLBN-2 |
| BchChl | Pseudooctadecabacter jejudonensis |
| BchChl | Pseudorhodobacter sp. MZDSW-24AT |
| BchChl | Pseudorhodoferax sp. Leaf267 |
| BchChl | Puniceibacterium antarcticum |
| BchChl | Rhizobacter sp. Root404 |
| BchChl | Rhizobiales bacterium |
| BchChl | Rhizobiales bacterium 65-79 |
| BchChl | Rhizobiales bacterium CCH10-E5 |
| BchChl | Rhizobiales bacterium KCTC 52945 |
| BchChl | Rhizobium ipomoeae |
| BchChl | Rhodobacter capsulatus |
| BchChl | Rhodobacteraceae bacterium |
| BchChl | Rhodobacteraceae bacterium CCMM004 |
| BchChl | Rhodobacteraceae bacterium CG17_big_fil_post_rev_8_21_14_2_50_63_15 |
| BchChl | Rhodobacteraceae bacterium EhC02 |

| | |
|---|---|
| BchChl | Rhodobacteraceae bacterium HLUCCA08 |
| BchChl | Rhodobacteraceae bacterium HLUCCA09 |
| BchChl | Rhodobacteraceae bacterium HLUCCO18 |
| BchChl | Rhodobacteraceae bacterium JBTF-M29 |
| BchChl | Rhodobacteraceae bacterium LMIT002 |
| BchChl | Rhodobacteraceae bacterium MA-7-27 |
| BchChl | Rhodobacteraceae bacterium O448 |
| BchChl | Rhodobacteraceae bacterium PARR1 |
| BchChl | Rhodobacteraceae bacterium SB2 |
| BchChl | Rhodobacteraceae bacterium SM1902 |
| BchChl | Rhodobacteraceae bacterium TG-679 |
| BchChl | Rhodobacteraceae bacterium THAF1 |
| BchChl | Rhodobacteraceae bacterium WDS1C4 |
| BchChl | Rhodobacteraceae bacterium WDS4C29 |
| BchChl | Rhodobacterales bacterium |
| BchChl | Rhodobium orientis |
| BchChl | Rhodoblastus acidophilus |
| BchChl | Rhodoferax antarcticus |
| BchChl | Rhodoligotrophos appendicifer |
| BchChl | Rhodomicrobium sp. JA980 |
| BchChl | Rhodopila globiformis |
| BchChl | Rhodoplanes piscinae |
| BchChl | Rhodopseudomonas palustris |
| BchChl | Rhodosalinus sediminis |
| BchChl | Rhodospira trueperi |
| BchChl | Rhodospirillaceae bacterium |
| BchChl | Rhodospirillaceae bacterium HHTR118 |
| BchChl | Rhodospirillales bacterium |
| BchChl | Rhodospirillales bacterium 20-60-12 |
| BchChl | Rhodospirillales bacterium 20-64-7 |
| BchChl | Rhodospirillum centenum |
| BchChl | Rhodothalassium salexigens |
| BchChl | Rhodovibrio salinarum |

| | |
|---|---|
| BchChl | Rhodovulum adriaticum |
| BchChl | Richelia intracellularis |
| BchChl | Rippkaea orientalis |
| BchChl | Rivibacter subsaxonicus |
| BchChl | Rivularia sp. PCC 7116 |
| BchChl | Roseateles depolymerans |
| BchChl | Roseiarcus fermentans |
| BchChl | Roseibaca calidilacus |
| BchChl | Roseibacterium elongatum |
| BchChl | Roseibium hamelinense |
| BchChl | Roseicitreum antarcticum |
| BchChl | Roseicyclus mahoneyensis |
| BchChl | Roseiflexus castenholzii |
| BchChl | Roseinatronobacter monicus |
| BchChl | Roseisalinus antarcticus |
| BchChl | Roseitalea porphyridii |
| BchChl | Roseivivax halodurans |
| BchChl | Roseobacter denitrificans |
| BchChl | Roseofilum reptotaenium AO1-A |
| BchChl | Roseomonas nepalensis |
| BchChl | Roseospirillum parvum |
| BchChl | Roseovarius aestuariivivens |
| BchChl | Rubidibacter lacunae |
| BchChl | Rubrimonas cliftonensis |
| BchChl | Rubritepida flocculans |
| BchChl | Rubrivivax benzoatilyticus |
| BchChl | Ruegeria sp. PBVC088 |
| BchChl | Salinarimonas rosea |
| BchChl | Salinisphaera sp. Q1T1-3 |
| BchChl | Salipiger mucosus |
| BchChl | Sandarakinorhabdus limnophila |
| BchChl | Scytonema millei |
| BchChl | Shimia sp. WX04 |

| | |
|---|---|
| BchChl | Skermanella aerolata |
| BchChl | Snowella sp. |
| BchChl | Sphaerospermopsis |
| BchChl | Sphingobium sp. TLA-22 |
| BchChl | Sphingomonas astaxanthinifaciens |
| BchChl | Spirulina major |
| BchChl | Stanieria sp. NIES-3757 |
| BchChl | Stappia sp. ES.058 |
| BchChl | Streptomyces purpurogeneiscleroticus |
| BchChl | Sulfitobacter guttiformis |
| BchChl | Synechococcus elongatus |
| BchChl | Synechocystis sp. PCC 7509 |
| BchChl | Tabrizicola aquatica |
| BchChl | Tardiphaga sp. vice154 |
| BchChl | Tateyamaria omphalii |
| BchChl | Telmatospirillum siberiense |
| BchChl | Tepidamorphus gemmatus |
| BchChl | Thalassobacter stenotrophicus |
| BchChl | Thalassobaculum litoreum |
| BchChl | Thalassobium sp. R2A62 |
| BchChl | Thalassococcus profundi |
| BchChl | Thermosynechococcus elongatus |
| BchChl | Thiobaca trueperi |
| BchChl | Thiocapsa marina |
| BchChl | Thiocystis violascens |
| BchChl | Thioflavicoccus mobilis |
| BchChl | Thiohalocapsa sp. ML1 |
| BchChl | Thiorhodococcus drewsii |
| BchChl | Thiorhodospira sibirica |
| BchChl | Thiorhodovibrio sp. 970 |
| BchChl | Tolypothrix campylonemoides |
| BchChl | Tranquillimonas alkanivorans |
| BchChl | Trichodesmium erythraeum |

| | |
|---|---|
| BchChl | Trichormus azollae |
| BchChl | Tropicibacter sp. LMIT003 |
| BchChl | Tropicimonas sp. IMCC34043 |
| BchChl | Tychonema bourrellyi |
| BchChl | unclassified Betaproteobacteria (miscellaneous) |
| BchChl | unclassified Rhizobium |
| BchChl | unclassified Sphingomonas |
| BchChl | uncultured marine proteobacterium |
| BchChl | uncultured marine type-A Synechococcus 4O4 |
| BchChl | uncultured marine type-A Synechococcus 5B2 |
| BchChl | uncultured marine type-A Synechococcus GOM 3O12 |
| BchChl | uncultured marine type-A Synechococcus GOM 5D20 |
| BchChl | uncultured proteobacterium DelRiverFos13D03 |
| BchChl | unicellular cyanobacterium SU2 |
| BchChl | Variovorax sp. KK3 |
| BchChl | Vulcanococcus limneticus |
| BchChl | Xenococcus sp. PCC 7305 |
| BchChl | Yoonia litorea |

**Table S3. Strains and plasmids used in this study.**

| Name | Type | Description | Source |
|---|---|---|---|
| DJ | Strain | Wild-type (WT); Nif+ | Dennis Dean, Virginia Tech |
| DJ2102 | Strain | Strep-tagged WT NifD; Nif+ | Dennis Dean, Virginia Tech |
| DJ2278 | Strain | Δ*nifD*::KanR; Nif- | Dennis Dean, Virginia Tech |
| DJ884 | Strain | *nifD*R187I mutant; Nif+(slow); overexpresses NifH | Dennis Dean, Virginia Tech |
| AK022* | Strain | Δ*nifHDK*::KanR; Nif- | This study |
| Anc1A ("AK013")* | Strain | Δ*nifD*::$nifD^{Anc1A}$; Nif+ | This study |
| Anc1B ("AK023")* | Strain | Δ*nifHDK*::$nifHDK^{Anc1B}$; Nif+ | This study |
| Anc2 ("AK014")* | Strain | Δ*nifD*::$nifD^{Anc2}$; Nif+ | |
| pAG25 | Plasmid | KanR cassette (APH(3')-I gene) + 400-bp *nifHDK* flanking homology regions, synthesized into XbaI/KpnI sites in pUC19; used to construct strain AK022 from DJ | This study |
| pAG13 | Plasmid | $nifD^{Anc1A}$ + 400-bp *nifD* flanking homology regions, synthesized into XbaI/KpnI sites in pUC19; used to construct strain Anc1A from AK022 | This study |
| pAG19 | Plasmid | $nifHDK^{Anc1B}$ + 400-bp *nifHDK* flanking homology regions, synthesized into XbaI/KpnI sites in pUC19; used to construct strain Anc1B from AK022 | This study |
| pAG14 | Plasmid | $nifD^{Anc2}$ + 400-bp *nifD* flanking homology regions, synthesized into XbaI/KpnI sites in pUC19; used to construct strain Anc2 from AK022 | This study |

*Kaçar Lab strain designations indicated for reference

**Table S4. Primers used in this study.**

| **Primer** | **Sequence (5' to 3')** | **Description** |
|---|---|---|
| 306_nifH_F | GCCGAACGTTCAAGTGGAAA | Forward primer, binds non-coding sequence upstream of *nifH; f*or PCR amplification of *nifHDK* and *nifH* sequencing |
| 307_nifH_R | AGAGCCAATCTGCCCTGTC | Reverse primer, binds non-coding sequence downstream of *nifH*; for *nifH* sequencing |
| 308_nifD_F | CACCCGTTACCCGCATATGA | Forward primer, binds non-coding sequence upstream of *nifD*; for *nifD* sequencing |
| 309_nifD_R | ACTCATCTGTGAACGGCGTT | Reverse primer, binds non-coding sequence downstream of *nifD*; for *nifD* sequencing |
| 310_nifK_F | GCTAACGCCGTTCACAGATG | Forward primer, binds non-coding sequence upstream of *nifK*; for *nifK* sequencing |
| 311_nifK_R | TCAGTTGGCCTTCGTCGTTG | Reverse primer, binds non-coding sequence downstream of *nifK*; for PCR amplification of *nifHDK* and *nifK* sequencing |
